# Divergent trajectories to structural diversity impact patient survival in high grade serous ovarian cancer

**DOI:** 10.1101/2024.01.12.575376

**Authors:** Ailith Ewing, Alison Meynert, Ryan Silk, Stuart Aitken, Devin P. Bendixsen, Michael Churchman, Stuart L. Brown, Alhafidz Hamdan, Joanne Mattocks, Graeme R. Grimes, Tracy Ballinger, Robert L. Hollis, C. Simon Herrington, John P. Thomson, Kitty Sherwood, Thomas Parry, Edward Esiri-Bloom, Clare Bartos, Ian Croy, Michelle Ferguson, Mairi Lennie, Trevor McGoldrick, Neil McPhail, Nadeem Siddiqui, Rosalind Glasspool, Melanie Mackean, Fiona Nussey, Brian McDade, Darren Ennis, The Scottish Genomes Partnership, Lynn McMahon, Athena Matakidou, Brian Dougherty, Ruth March, J. Carl Barrett, Iain A. McNeish, Andrew V. Biankin, Patricia Roxburgh, Charlie Gourley, Colin A. Semple

## Abstract

Deciphering the structural variation across tumour genomes is crucial to determine the events driving tumour progression and better understand tumour adaptation and evolution. High grade serous ovarian cancer (HGSOC) is an exemplar tumour type showing extreme, but poorly characterised structural diversity. We comprehensively describe the mutational landscape driving HGSOC, exploiting a large (N=324), deeply whole genome sequenced dataset. We reveal two divergent evolutionary trajectories, affecting patient survival and involving differing genomic environments. One involves homologous recombination repair deficiency (HRD) while the other is dominated by whole genome duplication (WGD) with frequent chromothripsis, breakage-fusion-bridges and extra-chromosomal DNA. These trajectories contribute to structural variation hotspots, containing novel candidate driver genes with significantly altered expression. While structural variation predominantly drives tumorigenesis, we also find high mtDNA mutation loads associated with shorter patient survival, and acting in combination with alterations in the nuclear genome to impact prognosis and suggesting new strategies for patient stratification.

## Introduction

High grade serous ovarian cancer (HGSOC) is the most common type of ovarian cancer. It usually presents with disease that has spread beyond the pelvis, and while initially sensitive to platinum-containing chemotherapy in 62% of cases, historically over 80% of cases relapse with a median overall survival of less than 5 years ^1,2^. The advent of first-line maintenance PARP inhibitor therapy is improving survival, particularly in patients whose tumours are homologous recombination repair deficient (HRD) ^3^. However, around 50% of tumours are not HRD ^4^ and the molecular drivers and therapeutic vulnerabilities in this patient subset with poorer prognosis are much less well characterised.

Recent studies of the HGSOC mutational landscape have noted the problems caused by structural complexity at many loci, potentially obscuring driver events and useful biomarkers^4^. The genomic aberrations at these repeatedly mutated loci require whole genome sequencing (WGS) data to be accurately resolved, and their combined impact determined. However, recurrent aberrations at a number of genes have been repeatedly reported. Initial analyses of the HGSOC genome using exome sequencing noted high levels of copy number alterations (CNAs) and predicted driver variants in a number of genes including: *TP53*, *NF1*, *RB1*, *CDK12*, *BRCA1*, and *BRCA2* ^5,6^. An early WGS study (N=80) recovered a similar list of recurrently disrupted genes, and also reported amplification of the *CCNE1* oncogene in 19% of samples ^7^. However, a more recent WGS study (N=118) identified less frequent driver alterations impacting additional genes, and reported recurrent deletions - rather than amplifications - at *CCNE1* in 45% of patients ^8^. The same study also suggested that the majority of driver events in HGSOC are likely to be mediated by somatic structural variants (SVs) and copy number alterations (CNAs). Thus, the driver landscape in HGSOC remains controversial, highlighting the need for comprehensive analyses of the mutational complexity of HGSOC genomes and elucidation of its clinical impact, in larger WGS cohorts.

HGSOC is subject to catastrophic mutational events, generating whole genome duplication (WGD) ^9,10^, complex structural variants (cSV) such as chromothripsis or ‘chromosome shattering’ ^11^, the production of extrachromosomal circular DNA (ecDNA) ^12^ and other cSV types involving overlapping amplifications and inversions ^13,14^. However, the interdependencies between cSV types have not been studied in detail in any tumour type, including HGSOC, and their individual and joint impacts on patient survival remain poorly understood ^15^. HGSOC tumour cells are also known to possess particularly abundant mitochondria carrying frequent somatic mtDNA mutations, though the best powered studies to date have failed to find associations between mitochondrial perturbation and patient outcomes ^16–18^.

Epistatic interactions between somatic mutations, across different scales of size and complexity, are thought to emerge frequently between different driver mutations during tumorigenesis ^19,20^, but remain poorly studied using WGS data. Several previous studies have used deep WGS to characterise HGSOC tumour samples ^5,7,21–23^ combined with gene expression profiling and other technologies, but have been constrained by modest sample sizes with limited power to detect recurrent mutations and any epistatic interactions between them.

Here we uniformly process previous WGS datasets combined with that from new patient samples to construct the largest deeply sequenced WGS combined cohort to date (N=324). We comprehensively describe the mutational landscape of HGSOC to define candidate driver genes subject to recurrent somatic single nucleotide variants (SNV), SV or CNA and for the first time determine the diversity and interactions of mutations driving tumorigenesis. We reveal the divergent evolutionary trajectories adopted by different HGSOC tumours to generate structural diversity, and gain new insights into the interdependencies of cSV events. We also perform the first comprehensive study of mtDNA mutations in HGSOC, revealing their association with patient survival. Finally, we construct an overarching model for HGSOC prognosis based upon all features of the mutational landscape and identify the most influential features.

## Methods

### Scottish sample collection and preparation for WGS and RNAseq

Scottish HGSOC samples (subsequently referred to as the SHGSOC cohort) were collected via local Bioresource facilities in Aberdeen, Dundee, Edinburgh and Glasgow as previously described ^22^. Clinical data for the SHGSOC cohort was retrieved from the Cancer Research UK Clinical Trials Unit Glasgow, the Edinburgh Ovarian Cancer Database and available electronic health records; the study received institutional review board approval from the Lothian Annotated Human BioResource (ethics reference 15/ES/0094-SR751) and NHS Greater Glasgow & Clyde Biorepository (ethics reference 22/WS/0020). HGSOC diagnosis was confirmed by formal expert pathology review (CSH) and samples were estimated to have >40% tumour cellularity by macroscopic visual assessment. Matched germline DNA was extracted from whole blood for each patient. Somatic and germline DNA was extracted from tumour and blood respectively as described previously ^22^. Somatic RNA was extracted from the same tumour sample as the DNA used for WGS. RNAseq was carried out by the Edinburgh Clinical Research Facility on an Illumina NExtSeq500 as previously described ^22^.

### Sequence acquisition and availability

WGS and RNA-seq reads were downloaded in compressed FASTQ format from the sequencing facility or in aligned BAM format (including unaligned reads) from the European Genome/Phenome Archive (Australian Ovarian Cancer Study (AOCS): EGAD00001000293, British Colombia Cancer Agency (BCCA): EGAD00001003268, MD Anderson (MDA): EGAD00001005240) and the Bionimbus Protected Data Cloud (The Cancer Genome Atlas (TCGA)). The reads obtained in BAM format were query-sorted and converted to FASTQ. The clinical information including survival end-points, age and FIGO (Fédération Internationale de Gynécologie et d’Obstétrique) stage at diagnosis is available for the AOCS and TCGA patients as part of the PCAWG project ^24^. Clinical information for the MDA and BCCA cohorts are available from the supplementary data of their respective publications ^21,23^. The SHGSOC cohort are available via EGA at accession number EGAS00001004410.

### Primary processing of RNA-seq

RNA-seq data was analysed using the Illumina RNA-seq best practice template. Briefly, reads were aligned to the hg38 reference genome and quality control was carried out. Salmon quant was used to quantify the expression of transcripts against the Ensembl 99 hg38 RefSeq transcript database indexed using the salmon index (k-mers of length 31). Transcript-level abundance estimates were imported into R and summarised for further gene-level analyses. For differential expression analyses, raw expression counts were used by the DESeq2 package. Previously published RNA-seq data available for the AOCS (N=80), TCGA (N=31) and MDA (N=26) cohorts, together with novel RNA-seq data for the SHGSOC (N=69) cohort, generated for the present study as detailed above, were processed in this way from FASTQ.

### Primary processing of WGS

Reads were aligned to the hg38 reference genome. Somatic and germline variant calling was performed using a bcbio ^25^ 1.0.7 pipeline as previously described ^22^. Germline SNPs and indels (SNVs) were called with GATK 4.0.0.0 HaplotypeCaller. Germline SNVs reported in ClinVar ^26^ to disrupt the function of 12 HGSOC risk genes ^27–29^ were recorded (Supp Table S2), and *BRCA1/2* variants were found to be enriched in patient samples relative to comparable populations in gnomAD ^30^. Somatic SNVs and indels were called as a majority vote between Mutect2 1.1.5, Strelka2 and VarDict 2017.11.23. Small variants were annotated with Ensembl Variant Effect Predictor v91 and filtered for oxidation artefacts by GATK 4.0.0.0 FilterByOrientationBias. Somatic structural variants were called with Manta 1.2.1 and GRIDSS 2.7.3. Somatic copy number alterations (CNAs) were called with CNVkit 0.9.2a0, CLImAT 1.2.2 and PURPLE 2.51. In both cases the intersect of the calls were taken forward as consensus calls. Consensus SVs were identified using viola-sv 1.0.0.dev10 ^31^ with a proximity threshold of 100bp. Consensus CNAs had at least 50% overlap between segments with the same direction of copy number change. Sample quality control was performed with Qsignature 0.1 to identify sample mix-ups and VerifyBamId 1.1.3 to identify sample contamination. Tumour cellularity was estimated using both CLImAT’s estimates and p53 variant allele frequency and CLImAT’s estimates were used throughout. To predict the level of HR deficiency in each tumour sample we implemented the HRDetect algorithm as published by Davies and Glodzik et al ^32^ as previously described ^22^.

### Clustering and classifying structural variation

We sought to determine whether a given SV belonged in a cluster of SVs or occurred alone. SVs occurring in clusters are expected to be more likely to represent the consequence of complex mutational processes such as punctuated catastrophic events. To cluster SVs we used the same approach as used by the PCAWG consortium (clusterSV ^33^). This approach identifies groups of SVs that occur closer together on a given chromosome than you would expect for that SV type on that chromosome given the overall distribution of breakpoints. To compare the SV landscapes between cohorts, we classified the consensus SVs. SVs that were not clustered were classified according to type, size, and reciprocity, in the case of inversions and translocations. This approach created 18 categories: clustered SVs; (simple) deletions (<100bp, 100 – 1kb, 1kb-10kb, 10kb-100kb, 100kb −1Mb, >=1Mb), duplications (<1kb, 1kb-10kb, 10kb-100kb, 100kb −1Mb, 1Mb – 10Mb, >=10Mb), inversions (large (>=1Mb) reciprocal, small (<1Mb) reciprocal, large (>=1Mb) unbalanced and small (>=1Mb) unbalanced), translocations (reciprocal or unbalanced). All size categories were verified using mixture modelling of size distributions for that SV type.

### Complex SV detection

Eight categories of cSV were predicted: chromothripsis, ecDNA, breakage fusion bridges, tyfonas, pyrgo, rigma, chromoplexy and seismic amplification. ShatterSeek^11^ was used to call regions subject to chromothripsis ^34^ based upon MANTA SV calls and CNVkit CNA calls, and following the recommended thresholds for the number of interleaved SVs, the number of adjacent segments oscillating between CN states, the number of interchromosomal translocations, the fragment joints test, the chromosomal enrichment test and the exponential distribution of breakpoints test ^11^. Candidate ecDNAs were predicted using AmpliconArchitect ^35^ with default settings to call ecDNA based upon purity and ploidy adjusted somatic CN calls from PURPLE. The resulting graph and cycle output files were processed by AmpliconClassifier^36^ to identify amplicons with CN>4 and size >10kb as ecDNAs. The junction-balanced genome graph (JaBbA) inference algorithm was used to generate genome graphs^13^. The resulting graphs were assessed as known cSV classes within gGnome ^37^, including breakage fusion bridges, tyfonas, pyrgo, rigma and chromoplexy. We detected putative regions of seismic amplification using an established approach ^14^ and the default threshold for amplification (CN >= 5 for diploid samples, and CN >= 9 for samples with ploidy >2). A candidate seismic amplicon was defined as one that contains amplified CN segments linked by >=14 SV rearrangements, as recommended^14^.

### CNA signature quantification

To robustly apply previously defined CN signatures ^38^ to higher resolution whole genome sequencing data, we filtered LogR and B allele frequencies from reads aligned to the whole genome to only include loci at Affymetrix Genome-Wide Human SNP Array 6.0 positions and recalled copy number using ASCAT 2.5.2. We then quantified the exposures of each of the copy number signatures in our samples using CINSignatureQuantification 1.1.2 with unrounded total copy number, as recommended ^38^. Individual signature exposures were compared between samples with and without *BRCA1/2* mutation using Wilcoxon rank-sum tests.

### Recurrently altered oncogenic pathways

We examined the extent that disruption via SNVs, CNAs and SVs was enriched within previously published oncogenic signalling pathways^39^. We examined ten canonical pathways: cell cycle, Hippo, Myc, Notch, Nrf2, PI-3-Kinase/Akt, RTK-RAS, TGFβ signalling, p53 and β-catenin/Wnt. The extent of enrichment of each type of alteration in each pathway was considered separately and represented by an oncoplot of pathway enrichment by variant type per sample using the R package complexHeatmap ^40^.

### Identification of genomic regions of recurrent copy number alteration/structural variation

Regions of the genome undergoing recurrent copy number alteration whether that be deletion or amplification were identified using GISTIC (v2.0.23). GISTIC compares the observed frequency of alteration in the region across all samples, combined with the observed magnitude of change, to the background expectation of frequency and magnitude of copy number alteration obtained by permuting genomic regions. We considered regions to be significantly recurrently deleted or amplified at a q-value < 0.05 and the region boundaries were defined to ensure 95% confidence that the reported region contained the recurrent event of interest. The CNA hotspot on chromosome X was identified by separate GISTIC analysis as GISTIC excludes chromosome X by default. GISTIC was applied to PURPLE segmentation calls only as these were more appropriately sized for use with GISTIC. More stringent filtering to consider only consensus calls within GISTIC peaks was applied downstream when investigating the impact of potential CNA drivers on expression. Broader chromosome-arm level recurrent gains and losses were also identified using GISTIC.

Genomic regions of significant (FDR corrected p-value < 0.05) SV enrichment were identified using a negative binomial regression model of SV density throughout the genome split into 50kb bins with 1kb overlap. All types of SV were considered separately in addition to breakpoint density. This model was implemented using the Fishhook package ^41^, adjusting for mappability (scores in 1Mb bins), GC content, CpG islands ^42^, gene density (over 5 Mbp windows; GENCODE v41), the presence of repetitive elements (DNA transposons, SINE, LINE, short tandem repeats, and long tandem repeats), and fragile sites ^43^. Genomic regions with low mappability including callable sites with mappability score < 0.9 ^44^ and blacklisted regions ^45^ were excluded. Chromosomal enrichments of SV classes were assessed using binomial tests of proportions accounting for chromosome length.

### Prediction of SNV mediated driver genes

We inferred SNV driver variants across the coding (canonical transcripts only) and non-coding genome separately for six genomic element types: promoter, 5’ UTR, coding, 3’ UTR, lncRNA and miRNA. We used four driver prediction methods: OncodriveCLUSTL ^46^, OncodriveFML ^47^, ActiveDriverWGS ^48^ and dNdScv ^49^, with differing approaches to driver identification. Prior to identification of driver variants, single nucleotide variants were filtered to remove variants with variant allele frequency (VAF) < 0.1. OncodriveCLUSTL v1.1.3 identifies unexpected clustering of SNVs. We optimized the OncodriveCLUSTL hyper-parameters (simulation window, smoothing window, and clustering window), selecting from a range of values for each, to maximise: the goodness of fit of observed p-values to the null distribution and the enrichment of known cancer genes in candidate driver elements. Otherwise we used default parameters and clusters were concatenated across genomic regions to identify candidate drivers at Q < 0.01. We used OncodriveFML v2.2.0 with default parameters to identify candidate variants based on bias in functional impact (Q < 0.01) and ActiveDriverWGS v1.2.0 to identify candidate variants based on mutational burden in functionally defined elements of interest (FDR < 0.05). Lastly, for coding regions only, we used dNdScv with default parameters to identify candidate variants based on the ratio of synonymous to non-synonymous mutations (qglobal cv < 0.01). Overall there was poor agreement between candidate drivers across algorithms (Supp Table S11). The results of dNdScv are expected to be most conservative, since it requires sufficient numbers of nonsynonymous SNVs per gene and uses a rigorous proxy for the background mutation rate. Six known HGSOC driver genes identified by dNdScv plus two additional novel candidate genes identified by dNdScv and at least one other algorithm were taken forward as candidate SNV mediated driver genes, and the full list of candidates from all algorithms is provided.

### Prediction of CNA/SV mediated driver genes

Genomic windows enriched for deletions, duplications, translocations, inversions and all breakpoints (at FDR < 0.05 by Fishhook) and CNA hotspots identified by GISTIC were intersected with COSMIC Cancer Gene Census (CGC) genes (version 96) and with the corresponding sample level consensus SV and CNA calls respectively. The CGC ^50^ is an established starting point for studies of cancer driver genes, containing 719 manually curated cancer driver genes, supported by functional validation studies and evidence of recurrent SNV/CNA in tumours. CGC genes in hotspots were tested for differential expression on occurrence of the SV/CNA type driving the hotspot using DESeq2. Samples in which the SV/CNA intersects the CGC gene were compared with those lacking an SV/CNA of that type in the gene being tested. Differential expression was tested separately in each of the four cohorts for which RNA seq data was available, and in a combined, batch corrected, table of expression values combining cohorts. An adjusted p-value of 0.05 for the gene being tested in the DESeq2 results was considered significant. The log2 fold change reported is that from the combined expression analysis, with the number of cohorts with a significant change reported as a replication score. An analysis of the FDR of this procedure suggested replication across cohorts maintained an FDR of less than 0.05.

### Analyses of mutual exclusivity and co-occurrence patterns across somatic alterations

Complex SVs were clustered according to their prevalence across samples using hierarchical clustering. The major axes of variation in the prevalence matrix (excluding HRD and WGD) were identified using principal components analyses and the first two components were compared in the presence of WGD and HRD. Patterns of pairwise co-occurrence and mutual exclusivity between complex SVs were explicitly tested based on weighted mutual information using SELECT ^51^ to extract significant pairwise relationships relative to expectation, whilst limiting the false discovery rate. This was repeated to consider all pairwise combinations of key nuclear somatic alterations, HRD and mtDNA mutations by mitochondrial complex. The fraction of tumour genome duplicated was compared in samples with and without breakage fusion bridges or chromothripsis.

### Calling mtDNA mutations and mtDNA copy number

For each sample, reads aligned to the mitochondrial reference genome were extracted from the alignment files using Samtools (v1.12). To mitigate against the issue of mismapped reads originating from nuclear-encoded mitochondrial pseudogenes (known as NUMTs), which could introduce false positives into the variant calling, the RtN! algorithm was used to filter out putative NUMT reads, including those not represented in the human reference genome, retaining only those from authentic mtDNA ^52^. The retained reads were then subjected to variant calling using VarScan2 (v2.4.4) ^53^ with the following parameters: -- strand-filter 1, --min-avg-qual 30 (minimum base quality 30), --min-coverage 2, --min-reads 2, and --min-var-freq 0. VarScan2 was selected as the preferred variant caller based on our own benchmarking analysis and its successful use in previous studies ^16,54^. To estimate the number of copies of mtDNA in tumour and normal samples, we applied a formula derived from the pan-cancer analysis of whole-genomes study (PCAWG)^16^.

To enhance the precision of our variant calls, we implemented a stringent set of filters. These filters excluded variants displaying significant strand bias (phred > 60), variants located within error-prone regions due to homopolymers (66-71, 300-316, 513-525, 3106-3107, 12418-12425, 16182-16194), variants with limited supporting reads (<30), and variants with extremely low heteroplasmy (<0.25% VAF). Subsequently, SNVs and INDELs were annotated using Ensembl’s Variant Effect Predictor (VEP) version 107, setting the -- distance parameter to 0. This annotation included pathogenicity predictions by SIFT ^55^ and PolyPhen2 ^56^. To validate the effective removal of reads originating from NUMTs, mtDNA variants were called in RNA-sequencing (RNA-seq) samples that were matched with a subset of the tumour WGS samples (206 out of 324 samples), since NUMTs lack evidence of transcription. Our analysis revealed that 88.3% of variants called in the WGS samples could also be identified in the corresponding RNA-seq data, and this percentage increased to 97.3% for variants above 10% heteroplasmy.

### Multivariable analyses of the impact of molecular factors on overall survival

The survival analyses considered 277 samples with complete overall survival time after diagnosis and tumour FIGO stage data. These samples were obtained from four cohorts (AOCS 80; BCCA 59; SHGSOC 110; TCGA 31). Overall survival times greater than 10 years were right censored (19 samples). The presence of: nine classes of complex SV (whole genome duplication, chromothripsis, pyrgo, chromoplexy, breakage fusion bridge, ecDNA, rigma, tyfonas, and seismic amplification); SNVs in eight candidate driver genes (*TP53, NF1, RB1, BRCA1, BRCA2, CDK12, SLC35G5* and *TAS2R43*); CNA or SVs in eleven candidate driver genes (*AXIN1, PCM1, WRN, FGFR1OP, LEPROTL1, DNMT3A, ARID1B, TCEA1, PTEN, CCNE1* (inversions and duplications), and *CEP89* (deletions and duplications)); and deleterious mitochondrial mutations were tested for their impact on survival. Overall survival was modelled using a Cox proportional hazards regression model using the coxph method in the R package survival. The effect of each variable was examined individually and in combination using a multivariable model adjusted for age (greater than mean age at diagnosis), FIGO stage (3 or above), HR deficiency as assessed by HRDetect, and stratified by cohort. Effect estimates with 95% confidence intervals were presented as forest plots. In addition, we applied cross-validated elastic-net regularised Cox regression to perform variable selection using the cv.glmnet method in the R package glmnet, revealing the most informative variables in the reduced forest plot.

## Results

### Extreme structural diversity dominates the HGSOC genome

The combined WGS cohort (N=324) constitutes the largest deeply sequenced (median 71X coverage; range: 52-136X; Supp Figure 1) primary HGSOC tumour dataset to date. WGS data from matched tumour and normal blood samples, and RNA-seq data from the same tumours, were uniformly remapped and analysed to generate a harmonised dataset of consensus somatic mutation calls and gene expression for five HGSOC WGS sub-cohorts: SHGSOC (N=115 ^22^), AOCS (N=80 ^7^), BCCA (N=59 ^21^), TCGA (N=44 ^5^), and MDA (N=26 ^23^). Although each sub-cohort was independently ascertained and sequenced, uniform processing and analysis revealed consistent mutational landscapes (Figure 1; Supp Figure 2). These landscapes are dominated by somatic SVs and CNAs occurring with similar rates and genomic spans (Supp Figure 2) rather than by SNVs (Supp Figure 3). SVs and CNAs are dominated by large duplications, with duplications composing 89% of all CNAs (Figure 1), consistent with previous studies ^33^. A variety of complex SVs (cSVs) are also highly prevalent in this tumour type (Figure 1A; Supp Table S1), particularly chromoplexy (55% of samples), chromothripsis (31%), pyrgo (28%) and breakage-fusion-bridge events (BFB) (27%). Diverse ecDNA species were observed in a minority (16%) of tumours across sub-cohorts and in 8% of tumours the predicted ecDNA structures contained an amplified oncogene (Supp Table S2). The only major difference in cSV occurrence between sub-cohorts was a significant enrichment (OR: 9.28; Chi-squared p=4.1×10^-13^) of ecDNA in AOCS samples, which are enriched for chemoresistant and relapsed tumours. As expected, samples also show enrichment of low-frequency germline variants known to increase the risk of HGSOC (Supp Table S3).

**Figure 1:**
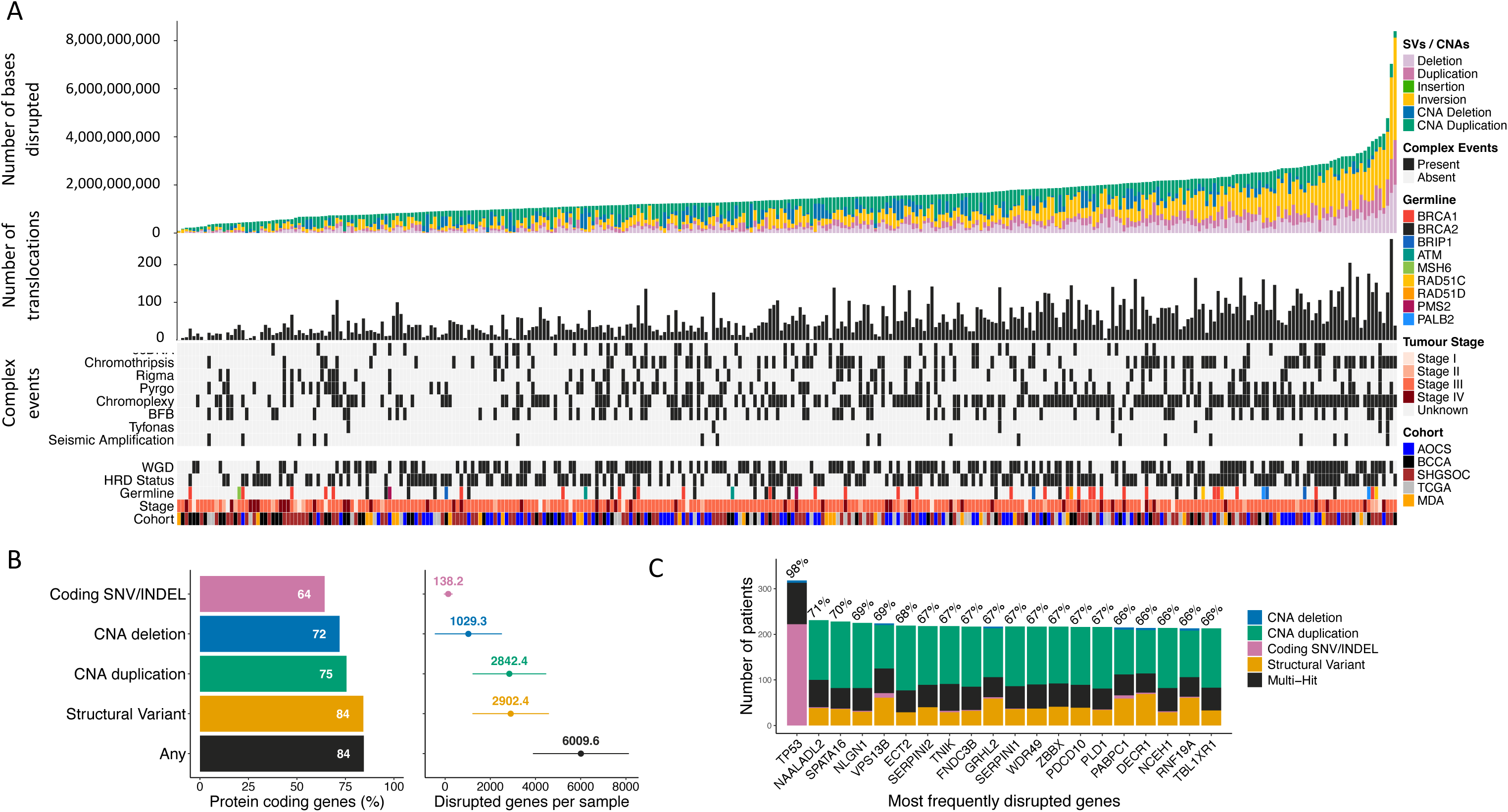
Structural complexity in HGSOC. (A) Structural variants are abundant across the combined cohort, reflected in the genomic span encompassed by SV and CNA calls and in the frequencies of translocations and complex events. Samples also show frequent HRD and WGD across tumour stages and cohorts. Pathogenic germline variants are relatively enriched in known HGSOC susceptibility genes. (B) Most protein-coding genes are disrupted by each class of variation across all samples (bar chart on left) but disrupted genes per sample (forest plot on right) are dominated by SVs and CNAs. Variants predicted to disrupt function are nonsynonymous SNVs of high/moderate impact and SVs or CNAs overlapping >=1 protein coding exon. (C) The most frequently disrupted genes overall are not enriched for known cancer genes, tend to be longer than average and to intersect recurrent CNA hotspots. The most frequently disrupted gene is TP53 where there is near ubiquitous mutation. NAALADL2 is an unusually long (1.37Mb) gene and is located within a common fragile site frequently altered across many tumour types (Li et al, 2020).

**Figure 2:**
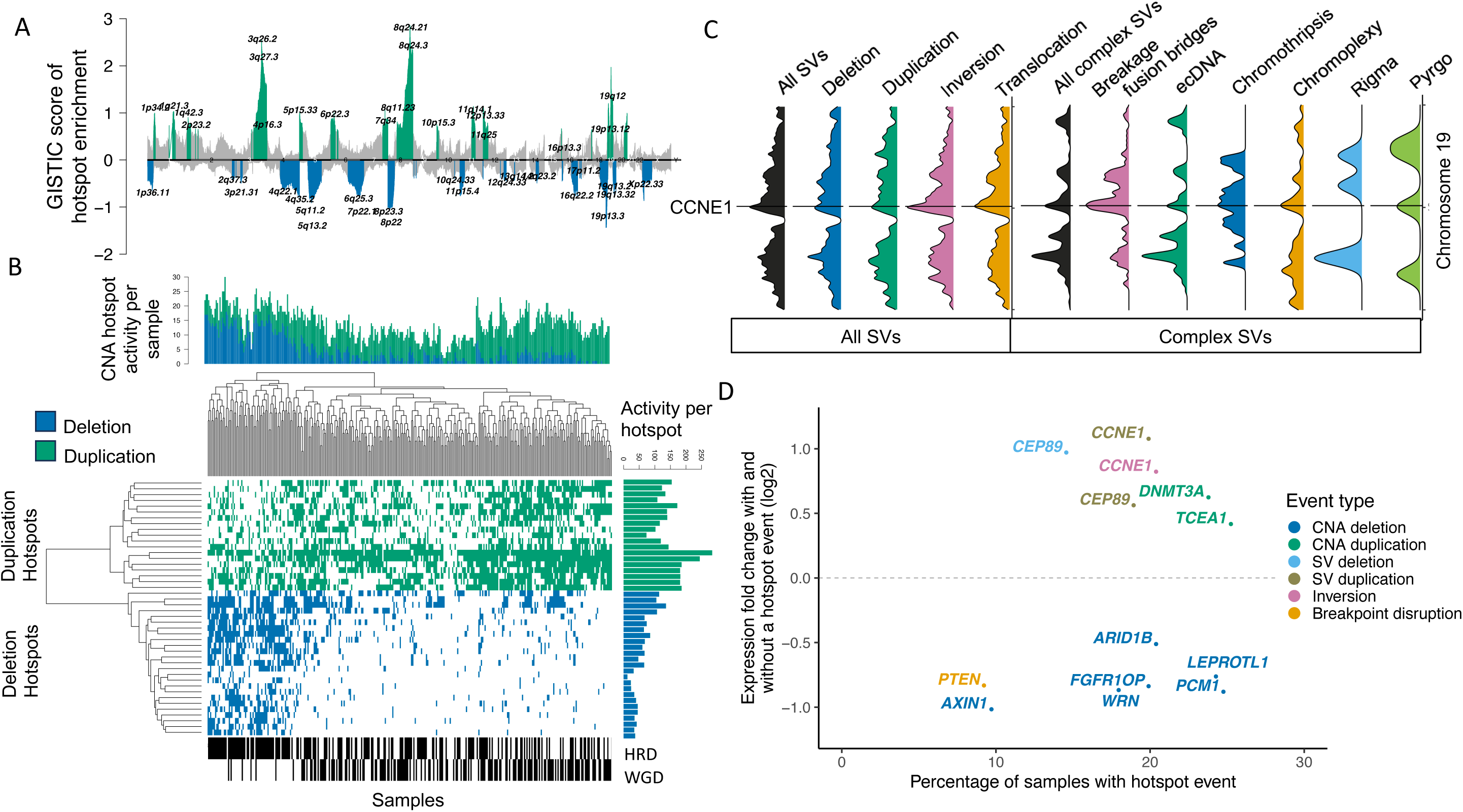
CNAs and SVs form hotspots throughout the genome which reveal candidate CNA/SV mediated driver genes. (A) GISTIC enrichment analysis reveals CNA hotspots. Duplication peaks are green and deletion peaks are blue. (B) CNA hotspot activity varies across samples with greater activity at deletion hotspots in HRD samples and greater duplication activity in WGD samples. (C) SVs are enriched on chromosome 19 and peak at CCNE1. In particular, there is a relative excess of inversions, translocations, breakage fusion bridges and chromothripsis implicating this gene. (D) Cancer gene census genes in SV/CNA hotspots that are differently expressed across samples in the presence of the event type driving the hotspot. Genes in CNA deletion hotspots or hotspots of breakpoint enrichment have significantly lower expression while genes in duplication (SV or CNA) or inversion hotspots have significantly higher expression. CEP89 SV deletions and SV duplications are both associated with increased expression reflecting the high level of overlapping genomic instability occurring at this locus which includes CCNE1. Expression fold changes are log2 transformed and are robust to the percentage of samples with the hotspot event.

**Figure 3:**
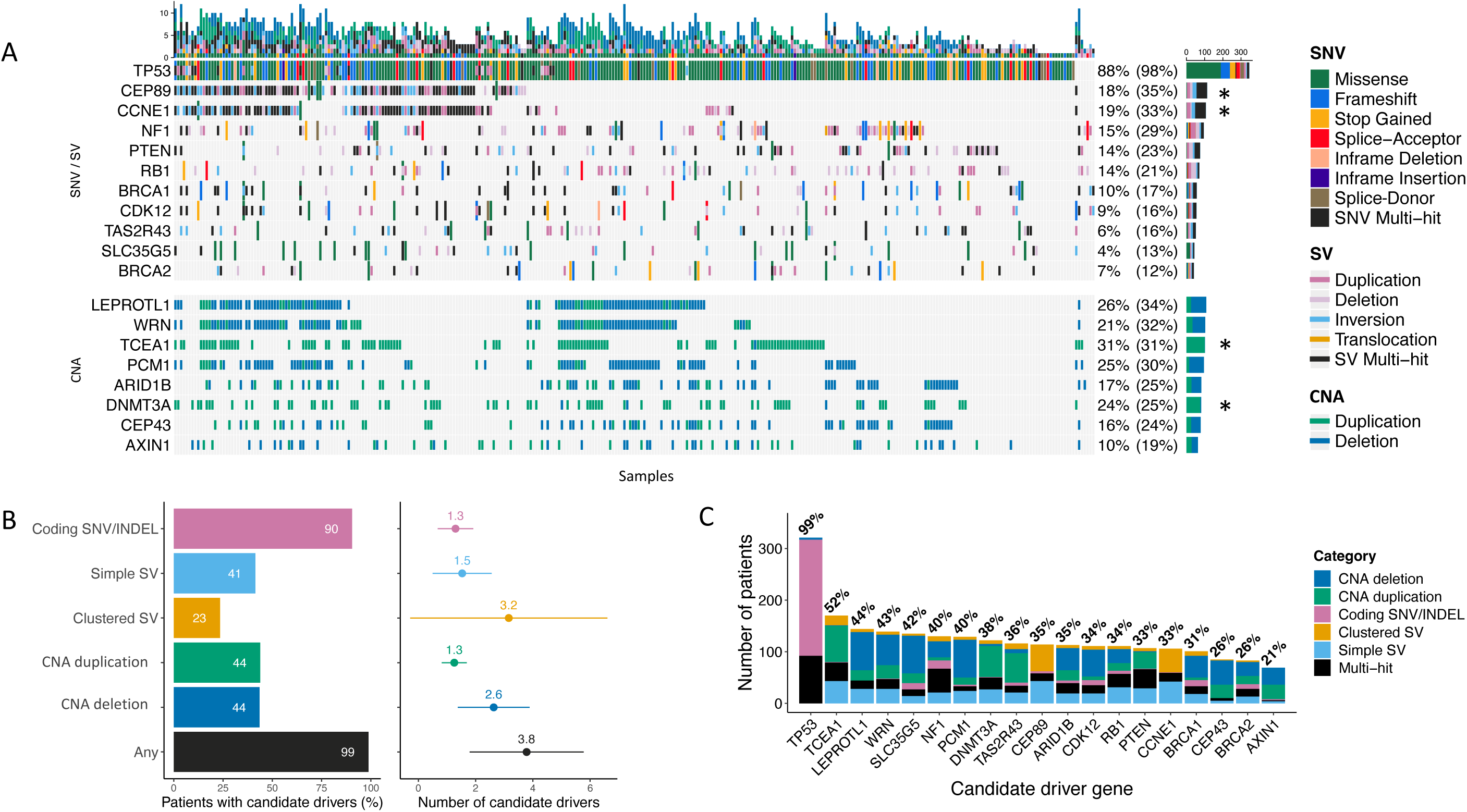
Diverse somatic mutation classes underlie HGSOC candidate driver genes. (A) Combined oncoplot indicates predicted driver genes (Methods) subject to recurrent pathogenic SNV/SV/CNA mutations predicted to impact function (SNVs annotated are nonsynonymous variants of HIGH or MODERATE impact by VEP; SVs or CNAs overlapping >=1 exon). The unbracketed percentage is the percentage of patients with a predicted pathogenic SNV/SV or CNA. For SNV/SVs this is either an SNV or a deletion, or in the case of the asterisked rows, representing genes associated with gain of function it is an SNV or duplication. The bracketed percentage is less conservative and is the percentage of patients with any annotated event. The vertical bar plot represents mutational burden of predicted deleterious SNVs/SVs.(B) The horizontal bar plot (left) represents the proportion of patients with different types of pathogenic driver mutations. The forest plot (right) represents the mean number of each type of driver mutation across tumours with at least one event and the standard deviation (whiskers), based on N=324 patients. (C) Total mutation frequencies in candidate driver genes. The proportions of patients with somatic alterations of any kind in each gene, whether predicted to be pathogenic or not. Clustered SVs are members of SV breakpoint clusters from ClusterSV (Li et al, 2020), the Multi-Hit category represents patients with >1 somatic alteration in the gene.

These events occur on a background of frequent HRD (56% of samples) and whole genome duplication (WGD; 49%). HRD and WGD are not mutually exclusive but are anti-correlated such that HRD is depleted in samples with WGD (odds ratio (OR) 0.56; Chi-squared p=0.015). HRD tumours are significantly enriched for deletions (Wilcoxon p-value <2.8×10^-5^) and WGD samples are significantly enriched for duplications (Wilcoxon p-value <1.8×10^-8^) as expected (Figure 2A, Supp Figure 3). The high frequency of CNAs provides insight into the underlying processes generating structural diversity, with dominant contributions of CNAs linked with HRD and chromosome mis-segregation, as reflected in the presence of known copy number signatures ^38^ as expected based on previous reports (Supp Figure 4; Supp Table S4).

**Figure 4:**
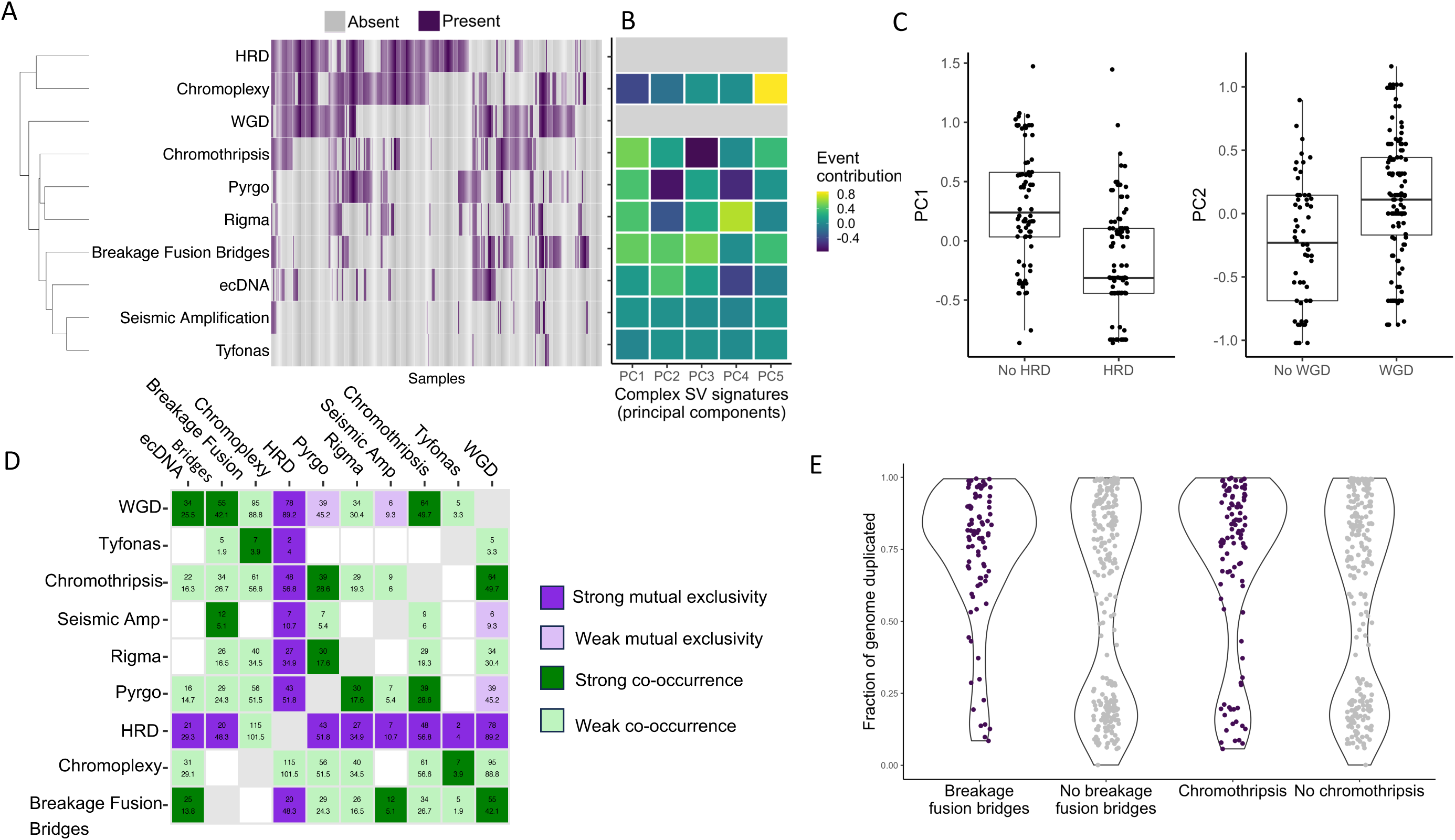
Patterns of co-occurrence and mutual exclusivity of complex structural variant classes and their relationship with WGD and HRD. (A) Abundance and co-occurrence of all complex events reveals two clusters of events defined by co-occurrence of HRD and chromoplexy, and co-occurrence of WGD and other cSV types. (B) Main axes of variation (principal components) in complex structural variant landscape across cohort. Heatmap represents feature loadings. HRD and WGD excluded from PCA. (C) HRD (PC1) and WGD (PC2) are associated with largest axes of variation in complex structural variant landscape across samples. PC1 significantly lower in HRD samples. PC2 significantly higher in WGD samples. (D) Biased co-occurrence and mutual exclusivity of cSV classes supports divergent tumour evolutionary trajectories with significant association seen between HRD and chromoplexy, but exclusivity between HRD and all other features. Cell counts represent observed vs expected counts of event co-occurrence in samples. (E) Association between the presence of chromothripsis and breakage fusion bridges (x-axis) and the estimated fraction of the genome duplicated (y-axis) across all samples (Bonferroni corrected chi-squared p for chromothripsis=6.7×10-3; breakage fusion bridges p=0.014).

The burden of SVs and CNAs per sample across the cohort are predicted to disrupt the function of several thousand genes in each tumour (Figure 1B), representing a large predicted deleterious mutation load. Many genes are impacted in the majority of samples, but with the exception of *TP53* they appear to be passenger rather than driver mutations (Figure 1C). For example, the top 10% of most heavily disrupted genes (N=1932) are only modestly enriched for genes with known roles in cancer (Cancer Gene Census (CGC)^50^)(OR: 1.5; Chi-squared p = 4.4×10^-4^). The majority of these genes appear to suffer collateral damage due to their proximity to SV/CNA hotspots. The compound phenotypic effects of high mutation rates at multiple levels of structural complexity on a given gene or pathway are challenging to predict. However, oncogenic signalling pathways ^39^ are likely to be disrupted in most samples, most prominently the RTK/RAS, NOTCH, Hippo and WNT pathways (Supp Figure 5). Intriguingly these pathways are often impacted by multiple pathogenic mutations, at multiple levels (SNV/SV/CNA) simultaneously, consistent with the presence of epistasis in tumour evolution, generating complex interactions between mutations in different genes in a pathway ^19^.

**Figure 5:**
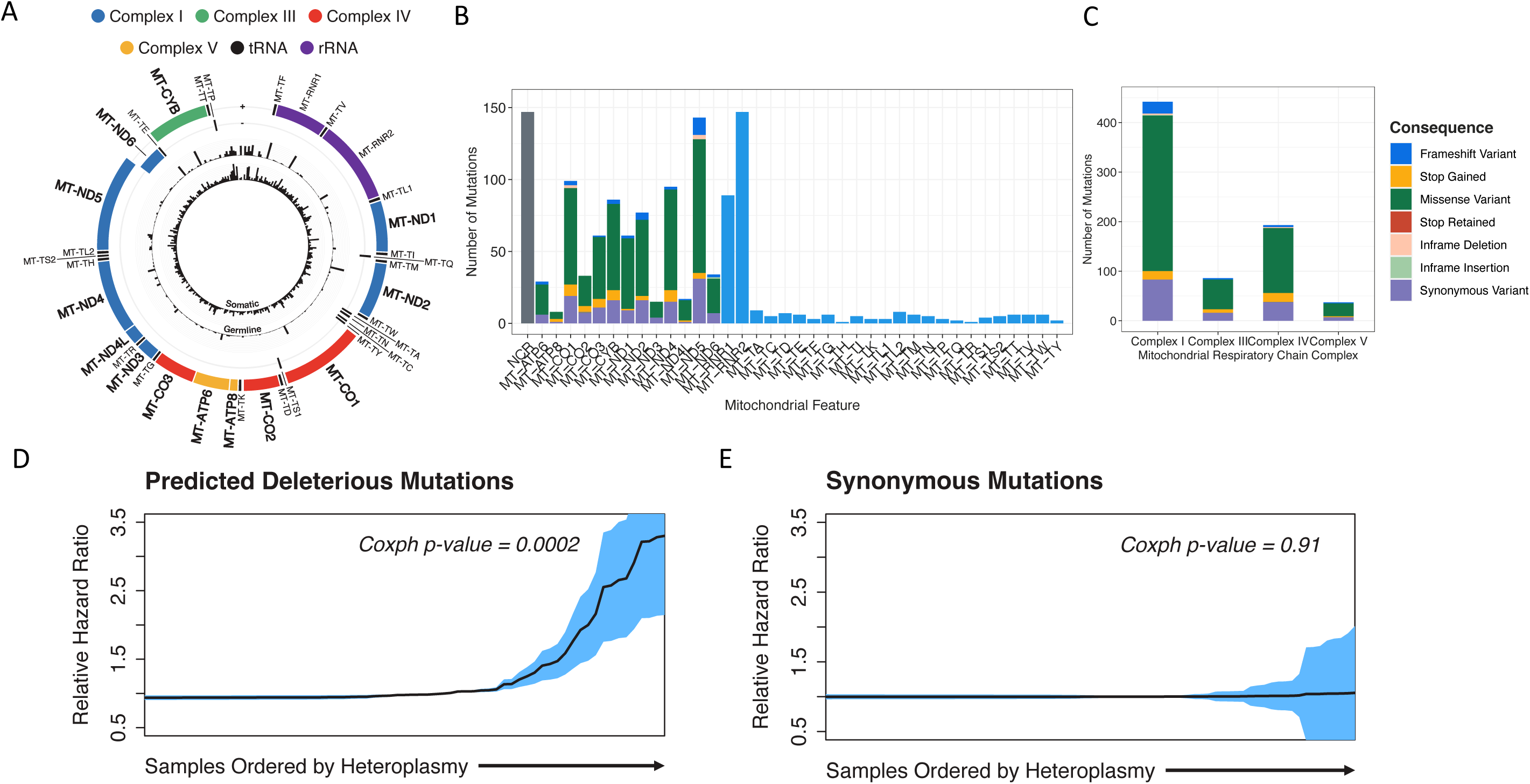
mtDNA mutation loads are a novel biomarker of overall survival. (A) Somatic (inner ring) and germline (outer) SNV frequencies across the cohort in mitochondrial encoded genes (black: single nucleotide variants; red: indels). (B) Abundant somatic SNVs disproportionately impact protein-coding genes in mtDNA. (C) SNVs categorized by VEP functional impact annotation include many protein altering variants expected to alter mitochondrial complex functions. (D) Overall survival Cox proportional hazards ratio increases with increasing heteroplasmy of deleterious mtDNA mutations (p-value=0.0002). (E) Overall survival Cox proportional hazards ratio is stable with increasing heteroplasmy of synonymous mtDNA mutations (p-value=0.91).

### Non-random distribution of complex structural variation across the genome

Both SVs and CNAs are non-randomly distributed across chromosomes in this cohort (Figure 2A; Supp Figures 6 and 7). We observed notable enrichments of all SV classes on chromosomes 8, 11, 12, 19 and 20 (Methods; Supp Figure 6A; Supp Table S5). Chromosome 19 also represents a novel hotspot for cSV in HGSOC as it is significantly enriched for BFB, chromothripsis, ecDNA and chromoplexy (Supp Figure 6B; Supp Table S5). Remarkably, this hotspot was seen across sub-cohorts, supporting the highly rearranged state of this chromosome as a general feature of HGSOC (Supp Table S5). The densities of SVs, chromothripsis and BFB on chromosome 19 peak at *CCNE1* (Figure 2C), a known HGSOC oncogene subject to recurrent amplification ^4,57^. Samples with BFB events at this locus also acquire higher *CCNE1* copy number amplifications than those with simple SV duplications (Supp Figure 8). *CCNE1* expression is also higher in the presence of SV duplications of any type and particularly so in the albeit few tumours where ecDNA and BFBs co-occur (Fold change relative to simple SV duplication: 2.8 (95% CI: 1.7 - 4.6), adj. p-value = 0.05) (Supp Table S6). This capability to amplify oncogenes beyond what is feasible via simple duplication events is likely of advantage to an evolving tumour. Similar effects as a result of ecDNA mediated amplification have been reported in other cancer types and are posited to lead to treatment resistance and subsequent poorer prognosis ^58^.

**Figure 6:**
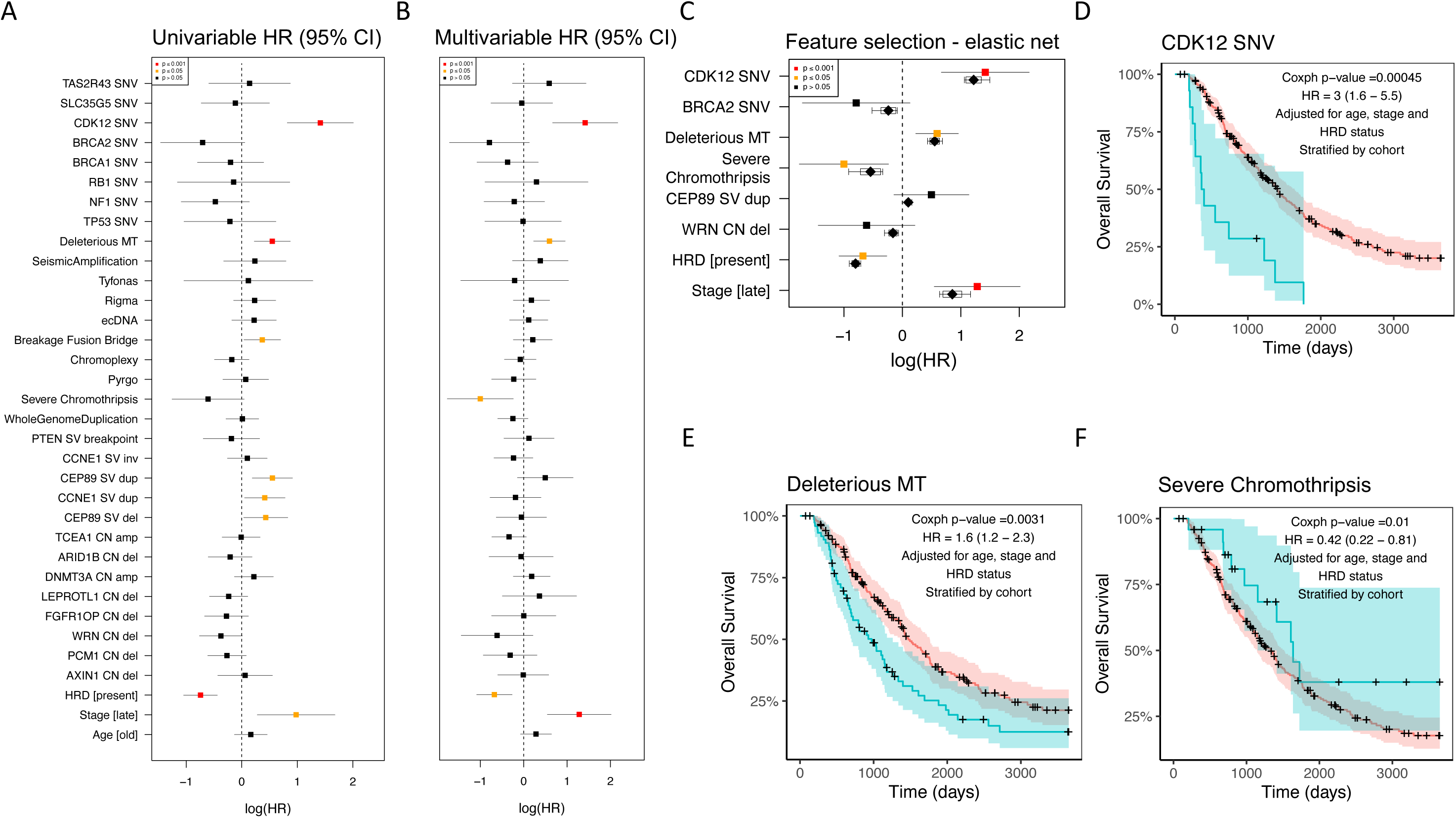
Multivariable modelling of the impact of genomic features (31) of HGSOC on overall survival adjusted for baseline clinical factors. (A) Univariable modelling of 31 genomic features using a Cox Proportional hazards model adjusted for HRD, age at diagnosis and stage at diagnosis and stratified by cohort. Forest plot shows hazards ratios (log) and 95% confidence interval per feature. B) As (A) but hazards ratios from one multivariable model including all 31 genomic features plus adjustments. (C) Selected features from elastic net penalised Cox PH regression model. Black diamonds are the median estimates from 10 cross-validations of elastic net. Kaplan-Meier curves of presence (blue curve) and absence (red curve) of (D) CDK12 SNV, (E) deleterious mtDNA mutation (F) severe chromothripsis.

We discovered additional genomic regions with significant enrichment of SVs - SV hotspots - at a number of loci across the genome (FDR < 0.05; Supp Table S7). In addition to the strongest signal across all SV types observed on chromosome 19, there are deletion and inversion hotspots on chromosome 2, a further inversion hotspot on chromosome 10 and multiple further loci enriched for SV breakpoints regardless of type throughout the genome. Moreover, we observe significant hotspots of breakpoints driven exclusively by translocations. We also determined CNA hotspots across the cohort based upon significant enrichment of variants in regions varying in size from tens of Kb up to multi-megabase regions encompassing entire chromosome arms (Methods; Supp Figure 9). Of 44 focal CNA hotspots examined, 25 were enriched for deletions and 19 for amplifications (Figure 2A Supp Table S8). The proportion of samples with CNAs at a given CNA hotspot varied from 4% to 42% (n=12 to 136 out of 324) and from 23% to 88% (n=73 to 284) for deletion and amplification hotspots respectively (Figure 2B). This suggests that even a relatively low level of recurrence, of deletions in particular, across samples is unlikely by chance and therefore informative. Chromosome arm-level events are frequent in HGSOC ^5^, in particular arm-level losses occur across many chromosomes in high proportions of samples (Supp Figure 9). Although CNA hotspot regions do include genes with known roles in cancer ^50^, they occur approximately in the numbers expected by chance (deletion hotspots OR = 1.2, CI: 0.88-1.7 Chi-squared p = 0.18; duplication hotspots OR = 1.4, CI: 0.99-1.96 p = 0.043). A critical question is therefore whether the expression patterns of such genes are altered, in response to the CNA burdens they incur, to affect the tumour phenotype.

### Hotspots of structural alteration implicate novel candidate driver genes

We adopted a novel approach to interrogate genes within CNA hotspots, exploiting the matched expression data available for the sub-cohorts making up the combined cohort to rigorously prioritise candidate drivers. We assume that CNA associated driver genes should show significant alterations in expression, consistent with the CNAs that impact them. For each cancer gene census gene present in a CNA hotspot (449 genes in total) we calculated the differential expression (DE) seen between samples with high copy number versus those with low copy number in each sub-cohort, and the associated false discovery rate (FDR) for DE genes seen across multiple sub-cohorts (Methods; Supp Table S9). Given the likely presence of confounding variation in the expression data (reflecting cellular heterogeneity and technical variation) these tests are necessarily conservative. Supporting this, we found that significantly lower expression of *PTEN* (a known tumour suppressor gene (TSG) in HGSOC) was associated with CNA deletion, but as this was seen in only one sub-cohort it failed to reach significance (FDR<0.05) and was excluded. Eight genes were identified as candidate drivers with several neighbouring genes altered simultaneously (Figure 2D). The expression of *AXIN1* (16p deletion, lower expression), *DNMT3A* (2q duplication, higher expression) and *TCEA1* (11q duplication, higher expression) all reflected the CNA burdens observed. Significantly lower expression of *PCM1, WRN* and *LEPROTL1* were associated with deletion, and all three are located within the same 8p deletion hotspot. Similarly, *ARID1B* and *FGFR1OP* are both within a 6q deletion hotspot, and both show lower expression in response to CNA deletions. Notably, the 8p deletion hotspot has been reported in many other tumour types ^59^ and may confer multiple advantageous traits on tumours ^60^. The genes underlying these effects were unknown, though recent work has shown that *WRN* deletion increases cell growth *in vitro*, suggesting *WRN* is a novel haploinsufficient pan-cancer TSG underlying the 8p deletion hotspot ^61^.

An analogous approach was taken to prioritise candidate genes based upon FDR corrected differential expression in SV hotspots (Figure 2D; Supp Table S10). Two genes emerged as SV associated driver candidates: significantly lower expression of *PTEN* (a known TSG in HGSOC) was associated with disruption by SV breakpoints, and higher expression of the HGSOC oncogene *CCNE1* was associated with both duplications and inversions (including foldback inversions where an inversion co-occurs with a duplication). The *CEP89* gene neighbouring *CCNE1* also passed the FDR threshold as a driver candidate, but was associated with both duplications and deletions, suggesting it may simply be a marker for the complex disruptions accumulating at the *CCNE1* locus rather than a driver in its own right. Genes and noncoding regions carrying recurrent SNVs were analysed using multiple driver prediction algorithms (Supp Figure 10; Supp Table S11) and recovered 6 known HGSOC driver genes (*TP53, NF1, BRCA1, CDK12, BRCA2, RB1*), plus another 2 genes with multiple paralogues: *SLC35G5* which has been reported previously as a source of artefacts ^62^ and *TAS2R43*. Many of the recurrently mutated genes have been identified as likely false positives in previous studies^63,64^ and in common with a recent study ^8^, we found no convincing evidence of SNV drivers in noncoding regions (Supp Figure 10).

The combined driver landscape - encompassing genes driven by SNVs, SVs and CNAs - is dominated by diverse structural alterations (Figure 3). The predicted pathogenic mutations (Figure 3A) reflect higher rates of alteration for known HGSOC genes than seen in previous studies lacking WGS data. As expected there appears to be improved diagnostic power for WGS in structurally diverse tumours relative to exome or panel sequencing. However it is possible that the higher rates seen in Figure 3A are underestimated, since the total mutation loads (regardless of pathogenicity) seen at driver genes are even higher (Figure 3C). For example, *NF1* pathogenic SNV/SV are seen in 15% of samples (Figure 1A), but in total 40% of samples show SNV/SV/CNA at *NF1* (Figure 1C). Disruption of *PTEN* and *RB1* have been reported as recurrent events in HGSOC ^4,65^ but the structural complexity seen at these loci ^7^ may have obscured inactivating alterations in studies lacking WGS data. Our current data confirm that these tumour suppressors are frequently disrupted by structural alterations, with pathogenic genomic events of any kind seen in 14% (*PTEN*) and 14% (*RB1*) of samples. The overall rates of alteration to *NF1*, *PTEN* and *RB1* are similar to those seen in another WGS project (N=118) ascertaining combined SNV/SV/CNA loads ^24^, which reported alteration rates of 24%, 14% and 19% respectively. Frequent pathogenic structural alterations to *BRCA1* and *BRCA2* are consistent with the role of SV/CNA mutations in HRD ^22^. Overall, each HGSOC sample is predicted to contain 3.8 driver variants on average with only 1.3 contributed by SNVs and the remainder involving SVs and CNAs (Figure 3B), which is similar to recent estimates based on WGS data (^24^; HGSOC N=118). In addition, many candidate driver SVs are members of significant SV clusters (Figure 3B, 3C) ^33^, which often indicate cSV events such as chromothripsis, suggesting that these events may be acting as drivers in some contexts. As we have shown tumours which are HRD or WGD differ in their genome-wide burden of deletion and duplication respectively, however, this appears to have no bearing on either the number or distribution of drivers by mutation type (Supp Figure 11).

### HRD and WGD underlie different evolutionary trajectories to structural diversity

The chaotic HGSOC genome harbours frequent occurrences of most cSV types reported to date, presenting a disordered and, at first sight, uninterpretable picture. Using state-of-the-art algorithms we have discovered high frequencies of known cSV types across the cohort, with chromoplexy, chromothripsis, pyrgo, rigma, BFB and ecDNA each seen in >10% of samples (Supp Table S1). This substantial WGS cohort is sufficiently powered to reveal significant biases in the patterns of cSV co-occurrence, a unique opportunity to study their genomic distributions and interactions in detail in patient samples, bringing clarity to our understanding of a tangled landscape.

We define two evolutionary trajectories to complex structural diversity (Figure 4). Each of these trajectories represent a major axis of variation in the cSV landscape in HGSOC, reflecting the underlying genomic state of the tumour. One trajectory involves HRD (Figure 4B, PC1) which is positively associated with chromoplexy, while the other involves WGD (Figure 4B, PC2) and a strong tendency to the acquisition of other cSV types (Figure 4B). Although these trajectories are not entirely mutually exclusive, it is evident that the divergent underlying genomic states of (i) deficiency in DNA repair and (ii) aneuploidy, are key aspects of HGSOC tumour biology which relate to different cSV profiles (Figure 4C). Striking patterns of mutual exclusivity exist between HRD and all cSV types except chromoplexy (Figure 4D). Particularly strong associations are seen between the fraction of the tumour genome duplicated and two highly disruptive cSV types - chromothripsis and BFB (Figure 4E). Abundant chromothripsis and BFB events account for disproportionate disruptions of genomic structure across the cohort, encompassing large fractions of the genome and causing many SV breakpoints in affected samples. The co-occurrence of these events with WGD suggests that WGD may buffer the particularly disruptive effects of these catastrophic events and limit their impacts on gene function.

Extensive characterisation of the HR proficient group of HGSOC is of great clinical importance as these patients have fewer options for targeted treatment, with patients with HRD tumours benefitting more from PARP inhibition. We observe that patients with a greater number of predicted chromothripsis events than average in their tumour genomes - or severe chromothripsis - had better prognoses than those patients with fewer or no chromothripsis events (Supp Table S12). Other cSV types, including ecDNA, showed weaker evidence for association with survival (Supp Table S12), contrary to previous pan-cancer reports ^15^ suggesting that the impact of ecDNA may differ in HGSOC from its impact in other tumour types.

### Mitochondrial and nuclear mutations combine to impact patient survival

Many tumour types, including HGSOC, are known to accumulate somatic SNVs in their mtDNA ^16^, but the consequences for mitochondrial (mt) function and patient survival remain unknown. We have found high mtDNA copy numbers and abundant somatic SNVs in tumour samples (Figure 5A; Supp Figure 12A; Supp Table S13), such that genes encoded in mtDNA suffer truncating and missense mutations at higher rates than in most other genes, including all known TSGs other than *TP53* (Supp Figures 12B and 12C). The highest deleterious mutation loads accumulate at particular genes (Figure 5B) and are predicted to disproportionately affect the function of mitochondrial Complex I (CI) and Complex IV (CIV) genes (Figure 5C). Remarkably, the predicted deleterious SNV loads in mtDNA are also a novel biomarker of poor patient prognosis, and mutations of higher heteroplasmy showing the largest effects (Figure 5D; Figure 6E). Notably, no associations with survival were seen for synonymous SNVs or SNVs occurring in mitochondrial RNA genes (Figure 5E; Supp Figures 13C and D), demonstrating that the impact on patient survival is mediated via the compromised functions of protein-coding mitochondrial genes, particularly those in CI.

Recent studies have reported deleterious somatic SNV loads in mitochondrial genes in renal, thyroid, and colorectal tumour types ^18^, but the associations of these loads with alterations to the nuclear genome are poorly studied. The rich mutational landscape of HGSOC described here provides an unusual opportunity to study these associations. Several trends emerge (Supp Figure 14) using a Bayesian inference approach ^20^ to study co-occurrence patterns across all somatic alterations. Firstly, within the mitochondrial genome there is an association between disrupted CI and CIII genes, suggesting specific alterations to mitochondrial metabolism. Secondly, these mitochondrial alterations significantly co-occur with WGD. Thirdly, this association appears to be attributable to WGD itself rather than the cSV (such as chromothripsis, BFB and ecDNA) that are correlated with WGD (Supp Fig 14A). In fact, the accumulation of somatic SNVs in most mitochondrial genes tends to be higher in the presence of WGD, and tends to be lower in tumours with HRD, suggesting less tolerance of disruptive mitochondrial DNA mutations in the presence of HRD (Supp Figure 14B). These interdependencies raise the question of whether the effects of mitochondrial mutation loads on survival are independent of the known influences of HRD, WGD and the many other somatic alterations of the nuclear genome.

We examined the dominant mutational features of the nuclear and mitochondrial genomes identified here to identify the variables driving differential survival. We systematically examined the associations of all individual features with overall survival (OS) time after diagnosis, including the presence of genomic aberration in genes with demonstrable recurrent SNV/SV/CNA (Figure 3), cSV types (Figure 4), and deleterious mitochondrial SNVs (Figure 5). Of these 34 binary features we found that 8 were individually associated with overall survival using Cox proportional hazards models stratified by cohort (Figure 6A; Supp Table S12). As expected, these 8 features included the presence of HRD and tumour FIGO stage, which have well established effects on OS. Other individually significant features were *CDK12* SNVs, deleterious mt SNVs, and the presence of severe chromothripsis. However, given the abundant interdependencies between these and other features (Supp Fig 14A) we employed integrative modelling to estimate the independent effects of all 34 features. Analysis using regularised Cox proportional hazards regression with an elastic net penalty, stratifying by cohort, revealed a refined model with redundant features pruned (Figure 6C; Supp Table S14). This model retained FIGO stage at diagnosis and HRD as well as 6 other features, of which 2 were significantly associated (adjusted p<0.05) in the multivariable model with overall survival. *CDK12* SNVs (Figure 6D) and deleterious mitochondrial SNVs (Figure 6E) were associated with poorer prognosis whereas the presence of severe chromothripsis (Figure 6F) was modestly associated with better prognosis given the available sample size and adjustment for multiple testing (Supp Table S14). *WRN* deletion, *CEP89* duplication at the *CCNE1* locus and *BRCA2* SNVs were also informative to the elastic net model although the evidence for their association with overall survival is limited in these data (Supp Figure 15).

## Discussion

We have shown that the global landscape of structural variation in HGSOC is shaped by the presence of HRD and WGD, leading to the emergence of hotspots, impacting thousands of genes recurrently across samples. Previous studies have identified similar genomic regions but have not refined these regions to identify candidate driver genes ^5,7,66^. Exploiting the independent sub-cohorts underlying the combined cohort, we predicted 8 novel HGSOC candidate driver genes (*PCM1, WRN, LEPROTL1, ARID1B, FGFR1OP, AXIN1, DNMT3A,TCEA1*) within these regions, showing significant differential expression across sub-cohorts in response to the CNA loads observed. Of these potential CNA-mediated driver events, only *WRN* (a DNA helicase involved in double strand break repair) deletion showed some evidence of an effect on patient survival. Supporting this conclusion, *WRN* has recently been reported as a haploinsufficient tumour suppressor gene, based upon analyses of pan-cancer CNA data not examined here and experiments in lung epithelial cells ^61^; we propose that it may also represent a novel therapeutic target in HGSOC.

Other CNA-mediated driver gene candidates provide new insights into HGSOC biology. Over-expression of the *de novo* methyltransferase *DNMT3A* has been reported in previous studies of HGSOC ^67^ and, consistent with this, we have shown that the frequent CNA amplification of this gene results in significantly higher expression in many samples. This suggests a role for *DNMT3A* in the aberrant DNA methylation patterns seen in HGSOC, which are not currently well understood, but show promise as biomarkers for detection and prognostic testing ^68^. *ARID1B* is a core subunit of the SWI/SNF chromatin remodelling complex and has been reported to be inactivated in endometrial, endometrioid ovarian and clear cell ovarian cancer ^69^, though inactivation of a similar subunit of the same complex (*ARID1A*) is more frequent. However, to our knowledge *ARID1B* has not previously been reported to be frequently altered in HGSOC. Activation of the Wnt/β-catenin pathway has long been reported in epithelial ovarian cancers, and it has been speculated that reductions in the expression of inhibitors of this pathway, such as *AXIN1*, could be a mechanism underlying pathway activation ^70^. We have found recurrent CNA significantly altering *AXIN1* expression, but intriguingly the rates of deletions and amplifications across samples are similar.

The extent of enrichment of SVs of all types, both simple and complex, on chromosome 19 is striking. To our knowledge this is the first report of a chromosome-wide hotspot for complex structural variation. This enrichment is driven by patterns of fold back inversions at the *CCNE1* locus ^21,57^ resulting in a higher level of amplification than possible by simple duplication and is achieved via complex mechanisms such as breakage fusion bridge cycles, ecDNA or chromothripsis. *CCNE1* amplification has been proposed as an effective therapeutic target ^71^ however the mechanisms leading to amplification and in particular over-expression of the gene are not fully understood. We show that SVs involving *CCNE1* are associated with its increased expression and our data suggest that concurrent duplication of nearby gene *CEP89*, which likely reflects the same amplification events, may be linked with poorer prognosis.

WGD is a common early event in many tumour types, including HGSOC, and has been associated with poor prognosis across cancer types ^9^. Our results confirm the association between WGD and genomic instability seen in cell line experiments ^72^, but extend this to encompass most known cSV types in addition to simple structural variation. It has been hypothesised that WGD may also allow rapid tumour evolution via catastrophic events such as chromothripsis ^73^, and we conclude there is convincing evidence for this in HGSOC. We have shown those tumours undergoing WGD suffer frequent catastrophic events, particularly chromothripsis and BFB, and would be expected to evolve rapidly. This may represent an advantage for some WGD tumours, for example those acquiring BFB mediated amplifications of *CCNE1*, but appears to be a liability for those suffering the most severe chromothripsis events, where patient survival is improved (Figure 6). This is consistent with simulations suggesting that WGD may be selected to mitigate the accumulation of deleterious alterations suffered by tumours with high mutation rates ^74^. A recent study found that HGSOC samples frequently showed evidence of chromothripsis but these events rarely caused losses of tumour suppressor or DNA repair genes ^11^. Our results also suggest that these events do not generally fuel adaptive evolution and instead make up part of the deleterious mutation burden afflicting these tumours. The presence of chromothripsis appears to be buffered by WGD and tumours with the most severe events may suffer increased immunogenicity or compromised metabolism, leading to longer OS (Figure 6).

Nevertheless, the frequent occurrence of WGD has clinical significance, since WGD itself has been reported to be a targetable vulnerability ^75,76^. This highlights the potential for new therapeutic opportunities in patients with WGD tumours which are generally HR proficient and currently more challenging to treat. Our knowledge of the cSV landscape is incomplete and the study of these variants is rapidly developing. Despite our increased power to characterise the cSV landscape in this larger cohort, the current known cSV types encompassed only a small fraction (13%) of the total SVs observed in the cohort. A higher proportion of SVs (27%) than those included in cSVs are clustered in the genome ^33^, which can indicate cSV. This suggests that as yet unstudied cSVs may occur in HGSOC.

We show that *CDK12* inactivation occurs at unexpectedly high levels in HGSOC, affecting up to 34% of samples in the cohort (Figure 3C) when all structural variation is added to the deleterious SNV/SV/CNA load affecting 16% of samples (Figure 3A). *CDK12* loss in HGSOC leads to genomic instability, in the form of extensive tandem duplication ^77^ and reduced expression of DNA damage repair genes including HR genes such as *BRCA1* ^78^. The extent to which CDK12 loss confers sensitivity to single agent PARP inhibition remains contentious; in prostate cancer the tandem duplication resulting from biallelic *CDK12* loss results in increased neoantigen generation and enhanced sensitivity to immunotherapy ^79^. We show that *CDK12* mutation is associated with significantly worse overall survival (Figure 6D) and it is possible that these patients, with genetic CDK12 inhibition, may show major improvements in overall survival with PARPi treatment.

Finally, we demonstrate that deleterious SNV loads predicted to disrupt mitochondrial gene function accumulate in HGSOC and are a novel biomarker of poorer OS, independent of other influential variables such as HRD. Recent studies have revealed driver roles for mtDNA mutations during tumorigenesis in particular tumour types, but the functional impact of these variants on mitochondrial function is under-studied ^18^. Notably, deleterious mtDNA mutations in colorectal tumours are associated with improved OS ^17^, highlighting the tumour type specific impacts of these mutations, and the pressing need for further studies. There are two clinical implications of our observations in HGSOC. Firstly, disrupted mitochondrial function may be a targetable feature, particularly in WGD tumours, and therapeutic strategies to rescue mitochondrial CI deficiency are already under investigation in the context of cardiovascular disease ^80^. Secondly, our data suggests that HRD tumours are intolerant of deleterious mtDNA mutations, consistent with their sensitivity to disrupted oxidative phosphorylation metabolism ^81^.

Overall, these data show that the genomic chaos seen in HGSOC obscures meaningful underlying patterns. Structural alterations are distributed non-randomly to generate hotspots harbouring known and novel driver genes. The diverse and frequent complex structural events observed relate to the presence of two genomic states, HRD and WGD, which generate structural diversity but also create vulnerabilities for tumours. Epitomising this dichotomy, tumours with WGD are more likely to possess *CCNE1* amplifications enhancing proliferation, but are also more likely to suffer extreme chromothripsis, which appears to impair tumour development. Thus, WGD tumours walk a narrow path towards an optimal level of chromosomal instability which facilitates rapid growth without risking cell death. These heavily disrupted nuclear genomes are in turn associated with alterations to the mitochondrial genome, impacting patient survival, and revealing a new layer of potential therapeutic targets.

## Acknowledgements

AE is supported by a University of Edinburgh Chancellor’s Fellowship and core funding from the MRC Human Genetics Unit. CAS and AM are supported by MRC core funding to the MRC Human Genetics Unit, University of Edinburgh and MRC Programme funding MC_UU_00035/1. EEB is an MB-PhD student supported by funding from CRUK (TRACC Programme SEBCATP-2022/100007). JT received core centre funding from CRUK. CG recieves research funding from AstraZeneca, MSD, Novartis, GSK, BerGen Bio, Medannex, Roche, Verastem, Artios and personal fees from AstraZeneca, MSD, GSK, Clovis, Verastem, Takeda, Eisai, Cor2Ed, Peer Voice. PR received research funding from the Beatson Cancer Charity and honoraria from AstraZeneca. RH received consultancy fees from GSK and DeciBio.

Sequencing of the SHGSOC cohort was supported by AstraZeneca, the Medical Research Council, and the Scottish Chief Scientist through a Precision Medicine Scotland Innovation Centre/Scottish Genome Partnership (SEHHD-CSO1175759/2158447) collaboration. This Scottish Genomes Partnership was funded by the Chief Scientist Office of the Scottish Government Health Directorates (SGP/1) and The Medical Research Council Whole Genome Sequencing for Health and Wealth Initiative (MC/PC/15080).This study would not be possible without the families, patients, clinicians, nurses, research scientists, laboratory staff, informaticians, and the wider Scottish Genomes Partnership team to whom we give grateful thanks. Members of the Scottish Genome Partnership (SGP) include Timothy J. Aitman, Andrew V. Biankin, Susanna L. Cooke, Wendy Inglis Humphrey, Sancha Martin, Lynne Mennie, Alison Meynert, Zosia Miedzybrodzka, Fiona Murphy, Craig Nourse, Javier Santoyo-Lopez, Colin A. Semple, and Nicola Williams. More information about SGP can be found at www.scottishgenomespartnership.org. The authors would also like to acknowledge the Edinburgh Clinical Research Facility for the sequencing of RNA samples from the SHGSOC cohort. The authors would also like to extend our thanks to the Nicola Murray Foundation and the Edinburgh Ovarian Cancer Database, from which the clinical data for much of the Scottish cohort were retrieved. We also thank the NRS Lothian Human Annotated Bioresource, NHS Lothian Department of Pathology, the Edinburgh Experimental Cancer Medicine Centre, the Biorepository at the Glasgow Queen Elizabeth University Hospital, and the Tayside Biorepository for their support. This manuscript was prepared using a limited access dataset obtained from BC CANCER and does not necessarily reflect the opinions or views of BC CANCER.

