## Supplementary Figures for "Divergent trajectories to structural diversity impact patient survival in high grade serous ovarian cancer"

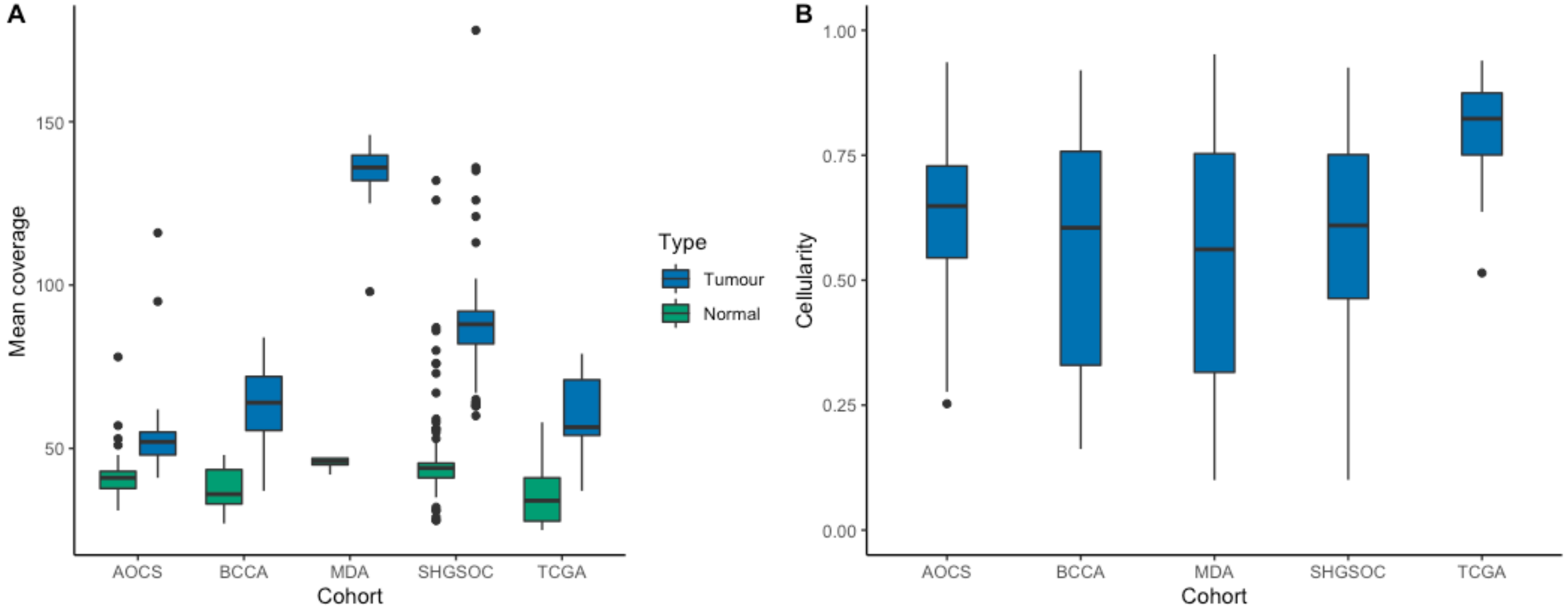


Supp Figure 1: Sequencing coverage and tumour purity across cohorts. (A) Mean coverage in tumour and normal samples across cohorts (AOCS: 52X, BCCA: 64X, MDA: 136X, SHGSOC: 88X, TCGA: 57X). Median coverage overall for the combined cohort was for tumours: 71X; and for normals: 42X. (B) Estimated purity across cohorts (overall median = 0.646, range 0.1-0.952).

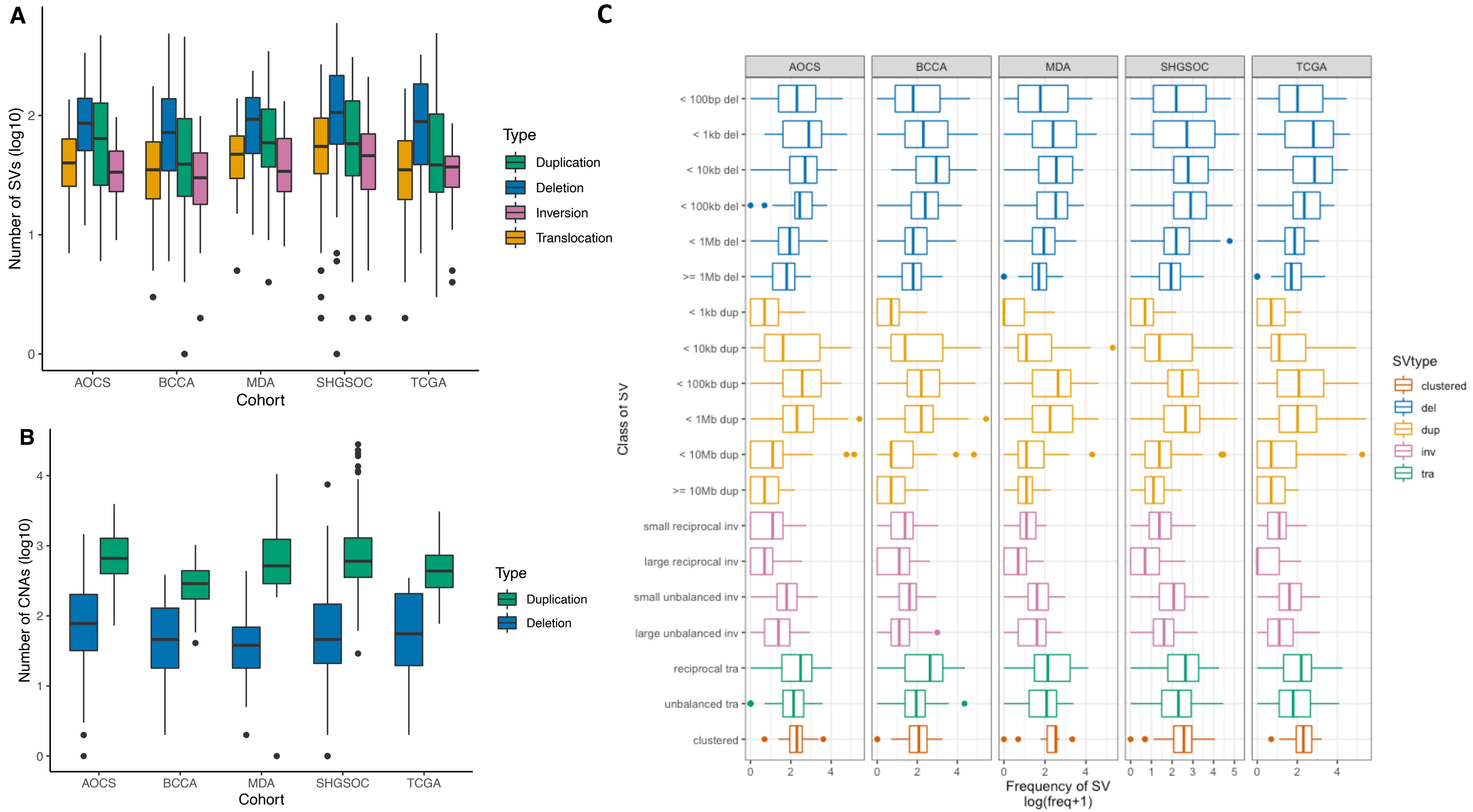


Supp Figure 2: SV and CNA burdens and genomic spans are broadly consistent across cohorts. (A) The numbers (log10) of consensus deletion, duplication, inversions and translocations by cohort. (B) The numbers (log10) of consensus deletion and duplication CNAs across cohorts. (C) The numbers of SVs by length and type across cohorts.

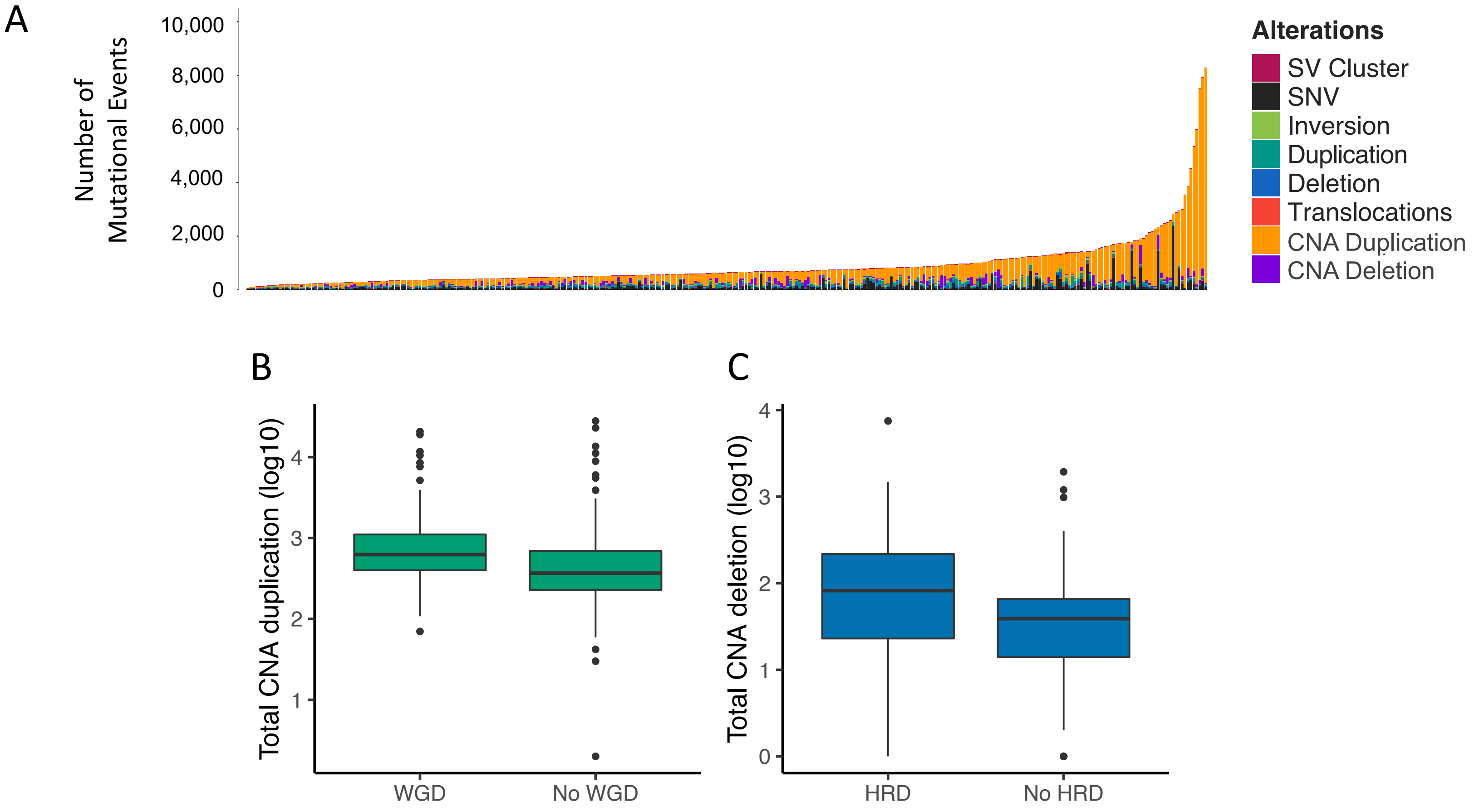


Supp Figure 3: SNV, SV and CNA burdens across cohorts. (A) The numbers of: SNVs, consensus SV deletion, duplication, inversions and translocations, consensus SVs in clusters and consensus CNAs across all samples. The mutational landscape is dominated by CNA duplications. (B) WGD samples have significantly higher duplication rates genome-wide per sample than samples lacking WGD (Wilcoxon $p<1.8x10^{-8}$); (C) HRD samples have significantly higher deletion rates than samples lacking HRD (Wilcoxon $p<2.8x10^{-5}$).

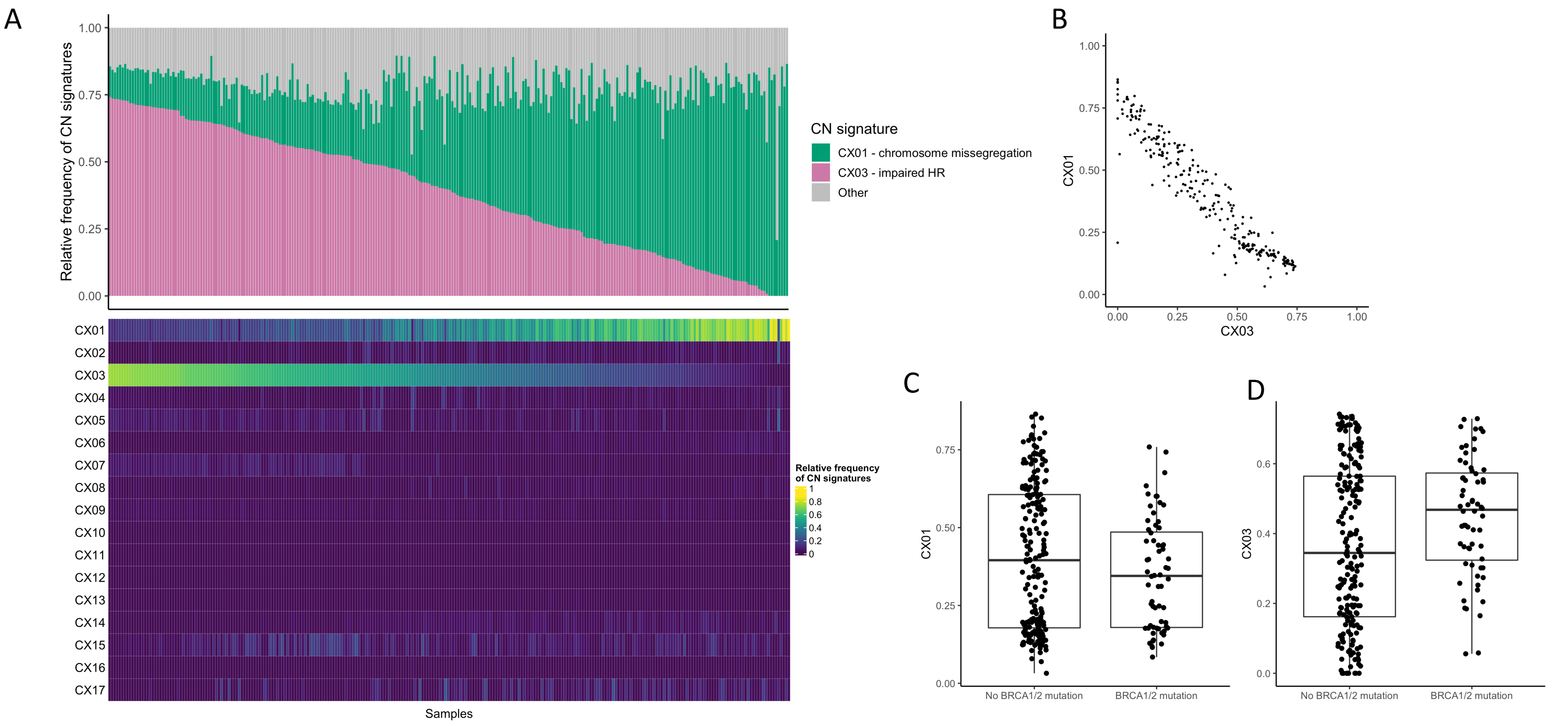


Supp Figure 4: CNA signature exposures reflect scars of impaired HR and chromosome missegregation. (A) Relative frequencies of exposures of known CN signatures (Drews et al, (2022)) across samples are dominated by signatures CX01 and CX03.(B) Exposures for CX01 and CX03 are strongly inversely correlated. (C) Exposures of CX01 in BRCA1/2 mutated and non-BRCA1/2 samples. (D) Exposures of CX03 in BRCA1/2 mutated and non-BRCA1/2 samples. CX03 exposure significantly higher in BRCA1/2 mutated samples (p=0.01). Only samples with > 40% cellularity (n = 268) were included as recommended by Drews et al.

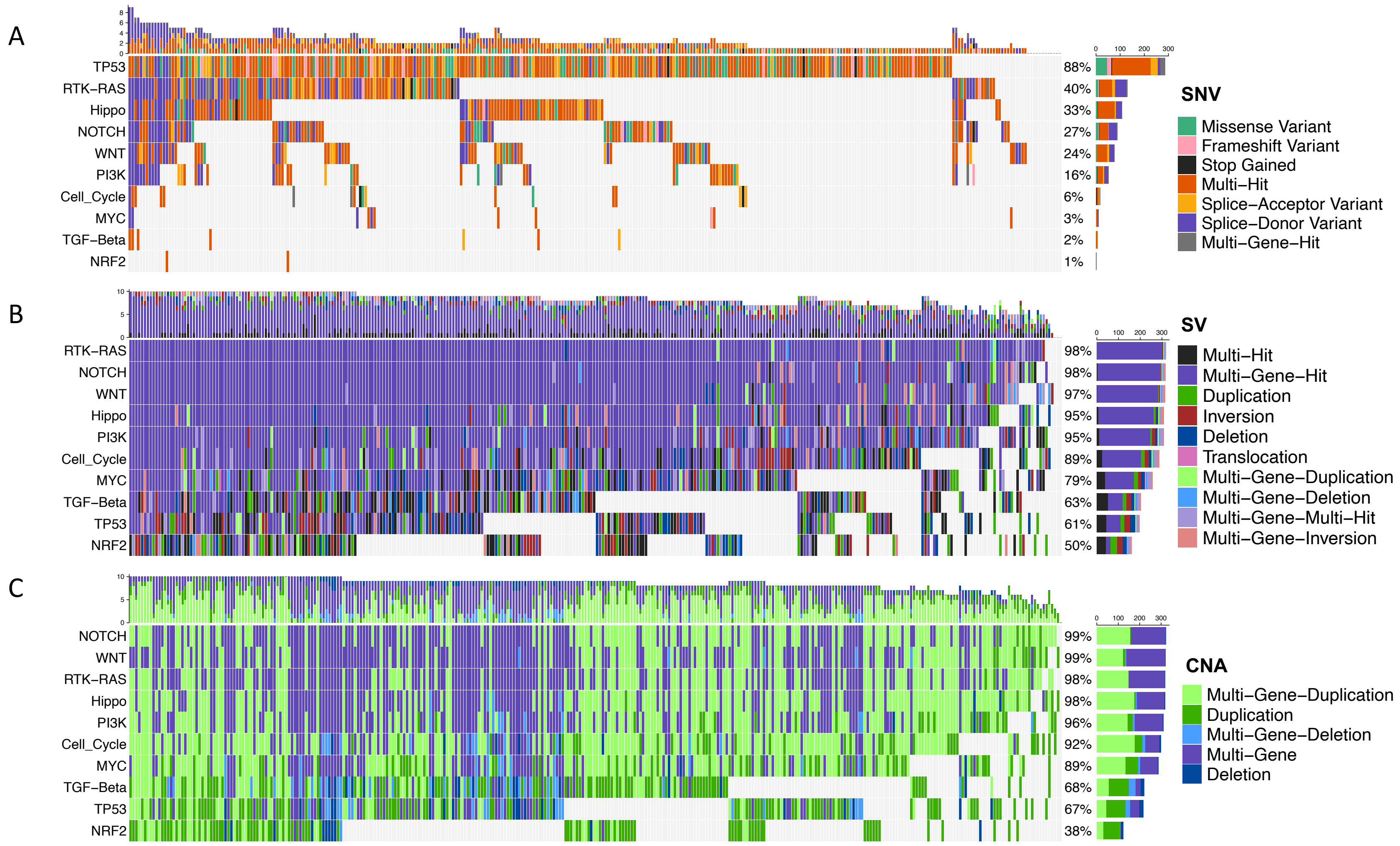


Supp Figure 5: Recurrently altered oncogenic pathways. (A) SNV oncoplot showing samples (x-axis) with oncogenic signalling pathways (y-axis; Sanchez-Vega et al, 2018) recurrently disrupted by SNVs across the cohort. Samples indicated have one or more predicted deleterious mutations (missense variants predicted by SIFT+PolyPhen) in a pathway associated gene. (B) SV oncoplot showing samples (x-axis) with pathways (Sanchez-Vega et al, 2018) where one or more SV intersects one or more exons of a pathway associated gene. (C) CNA oncoplot showing samples (x-axis) with pathways (Sanchez-Vega et al, 2018) where one or more CNA intersects one or more exons of a pathway associated gene. Single-Gene Multi-Hit = a single gene within the pathway carries multiple mutations. Multi-Gene-Hit = multiple genes within the pathway are disrupted by different SV/CNA types. Multi-Gene Multi-Hit = multiple genes within the pathway are disrupted by different SV types. Multi-Gene SV = multiple genes within the pathway are disrupted by the same SV type.

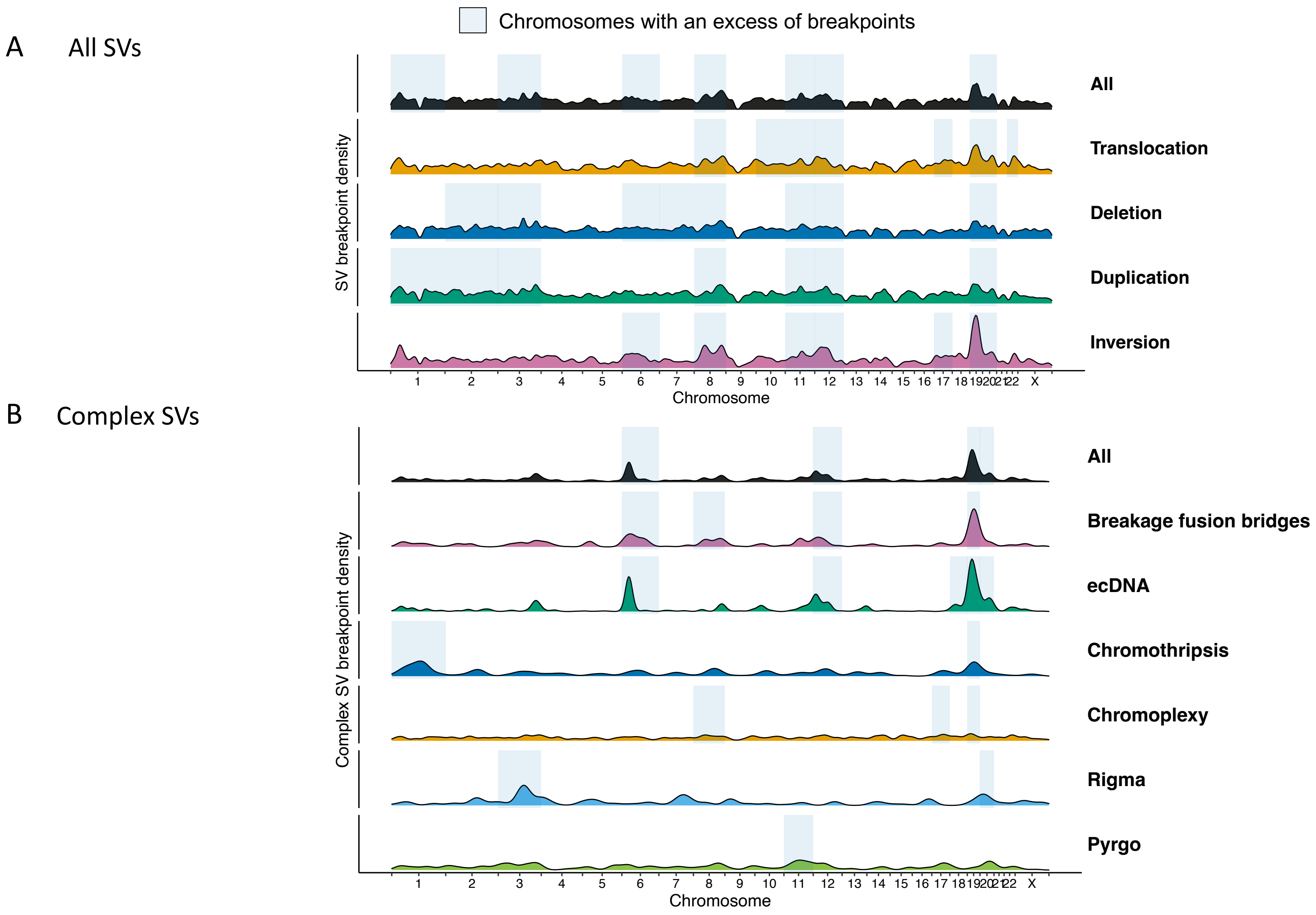


Supp Figure 6: Chromosome-level enrichment of breakpoints associated with SVs and complex SVs across the genome. (A) SV breakpoint density throughout the genome for all classes of SV. (B) Breakpoint density of SVs associated with complex SVs throughout the genome by complex SV type. Highlighted chromosomes have a significant excess of breakpoints adjusted for chromosome size (binomial test of proportions, $p<0.05$). Y-axis scales are consistent across SV types within each panel.

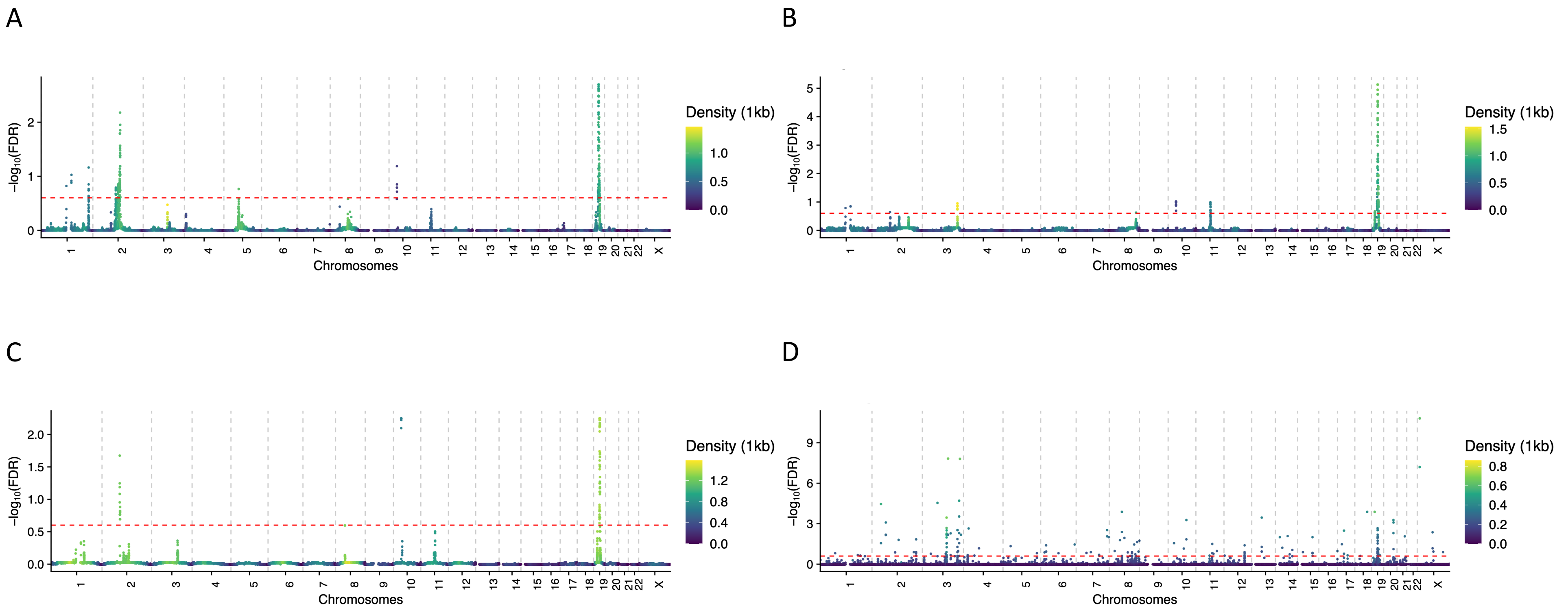


Supp Figure 7: Recurrently SV altered genomic regions. Genomic regions recurrently impacted by individual SV types across the cohort based upon FishHook analysis of consensus SV data in 50Kb windows, for (A) recurrent deletions, (B) duplications, (C) inversions and (D) breakpoints. The red lines indicate the FDR < 0.25 threshold which is far lower than the height of the peaks.

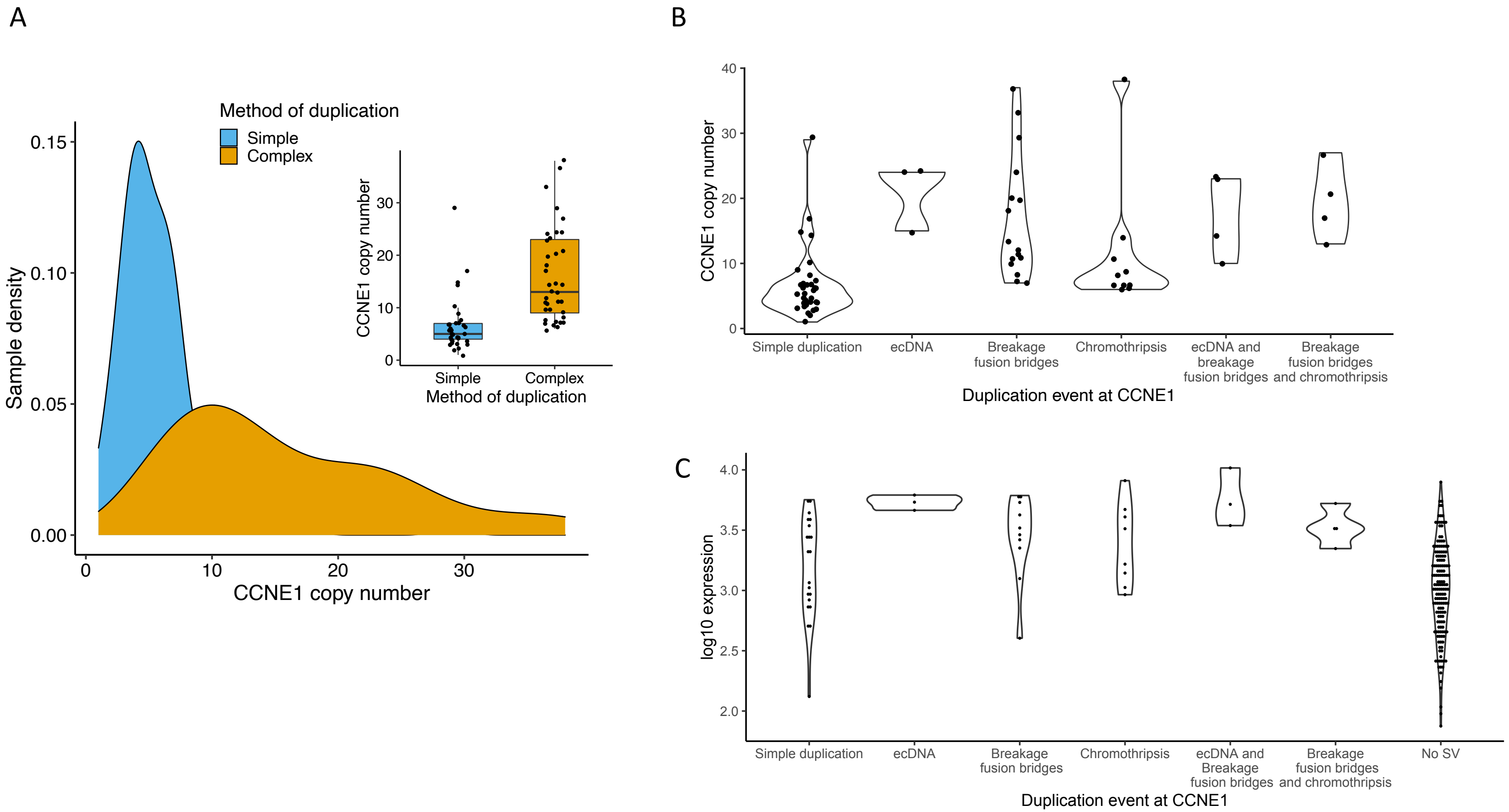


Supp Figure 8: Amplification of CCNE1 by simple SVs and cSV types. (A) The copy number of CCNE1 amplified by complex SVs or simple duplication differs significantly (Wilcoxon $p = 5.24x10-8$). CCNE1 is amplified to a large range of higher copy numbers by complex SV types. Amplification by simple SV is generally to a smaller range of lower copy numbers. (B) Copy number of CCNE1 amplified by simple SV events and observed combinations of cSV. All combinations of complex SV mediated amplification are associated with a higher degree of amplification.

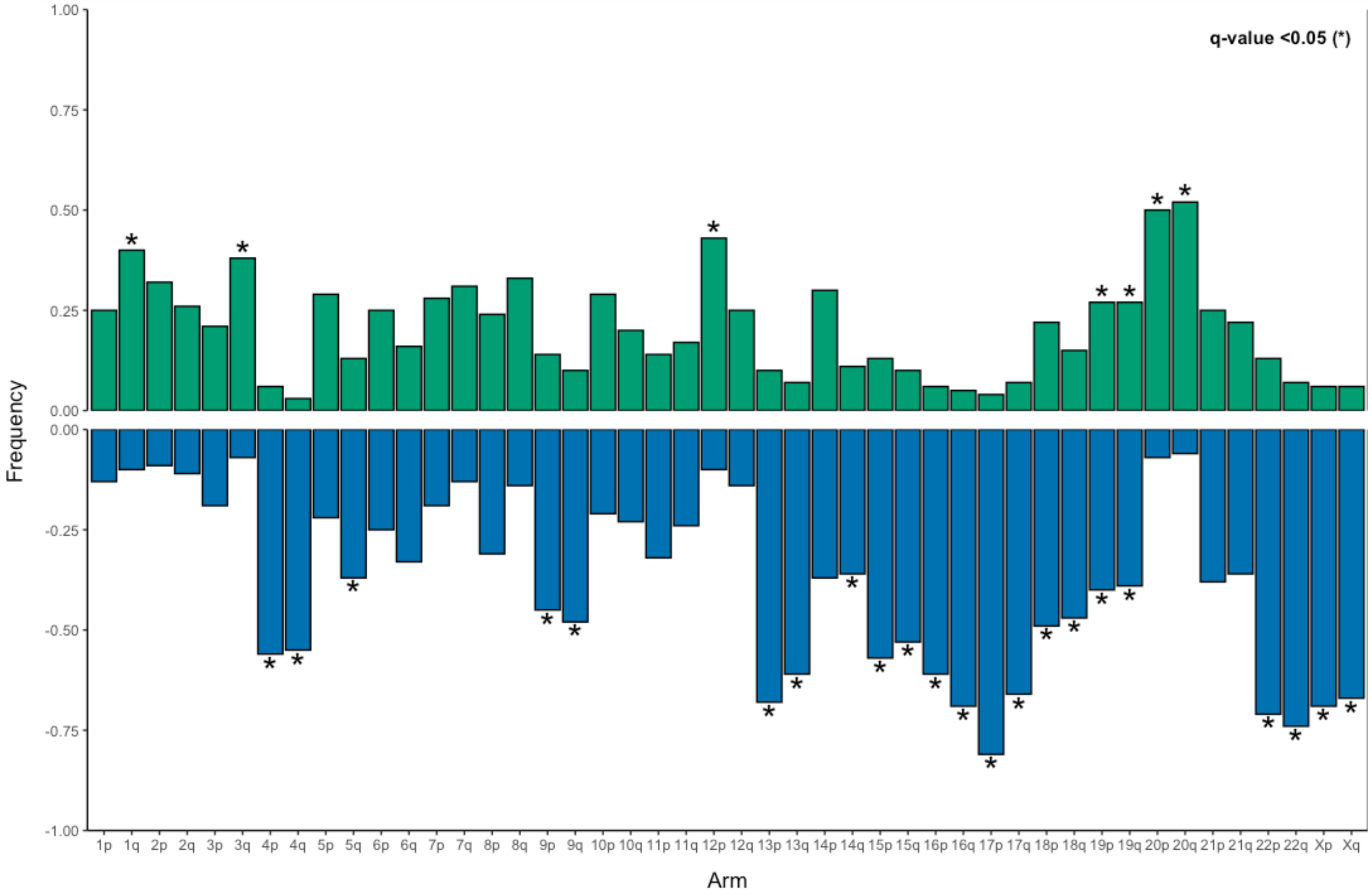


Supp Figure 9: Frequency of chromosome arm-level CNA across the genome. Chromosome arm-level alterations showing significant recurrence are indicated by asterisks. Arm-level losses are in blue and arm-level gains are in green.

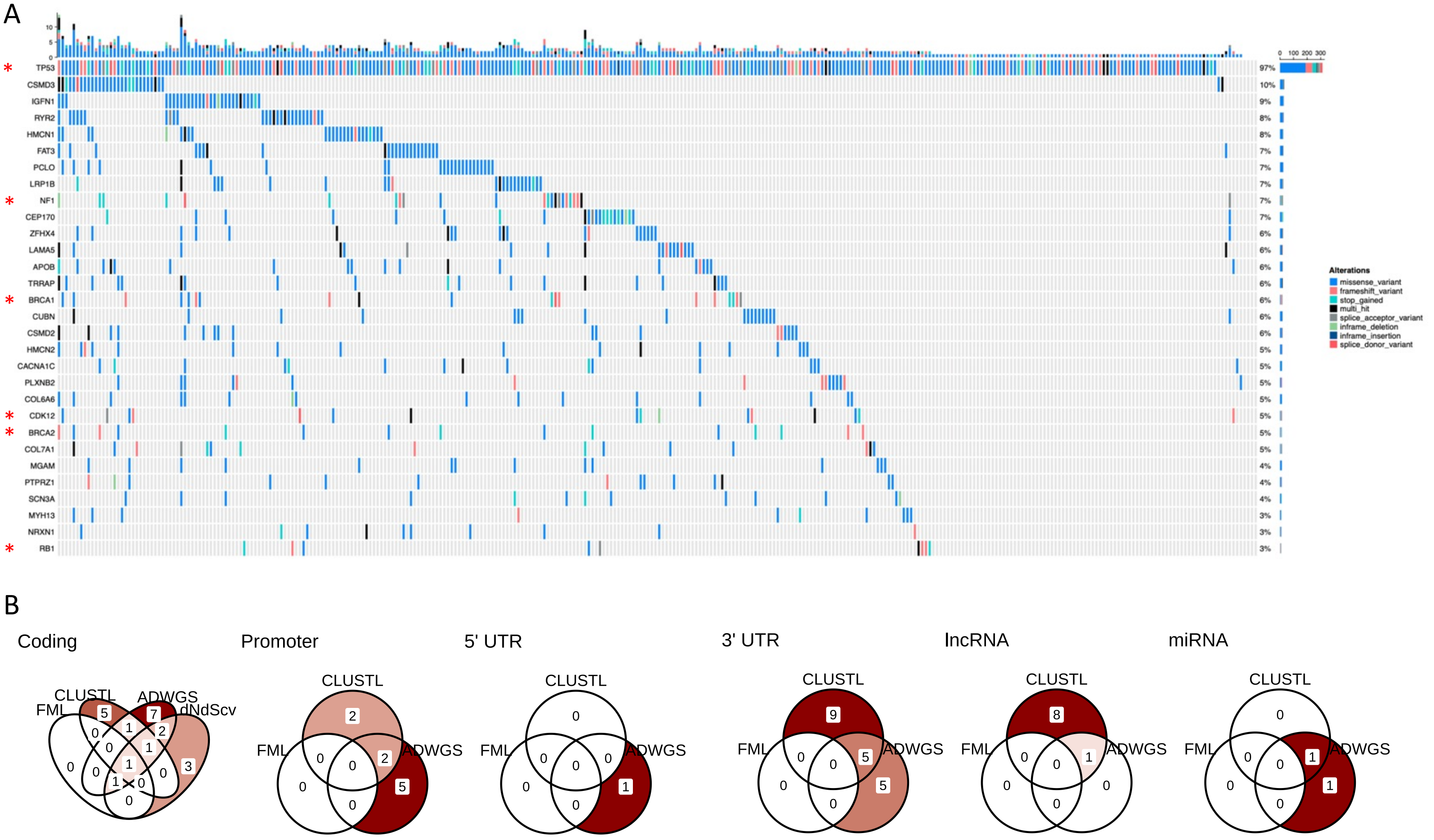


Supp Figure 10: Recurrently SNV altered genes and candidate driver variants. (A) SNV oncoplot (corrected for gene length) indicating protein coding genes subject to recurrent SNVs across the cohort and predicted to be driver genes (red stars) by dNdSCV. Many of these recurrently mutated genes have been identified as likely false positives as they are generally more frequently mutated (FLAGS: https://bmcmedgenomics.biomedcentral.com/articles/10.1186/s12920-014-0064-y, MutSigCV: https://pubmed.ncbi.nlm.nih.gov/23770567/). (B) Consensus driver prediction across four algorithms (dNdSCV, oncodriveCLUSTL, oncodriveFML and ActiveDriverWGS) indicating a lack of convincing evidence for novel SNV driver variants in coding or noncoding regions.

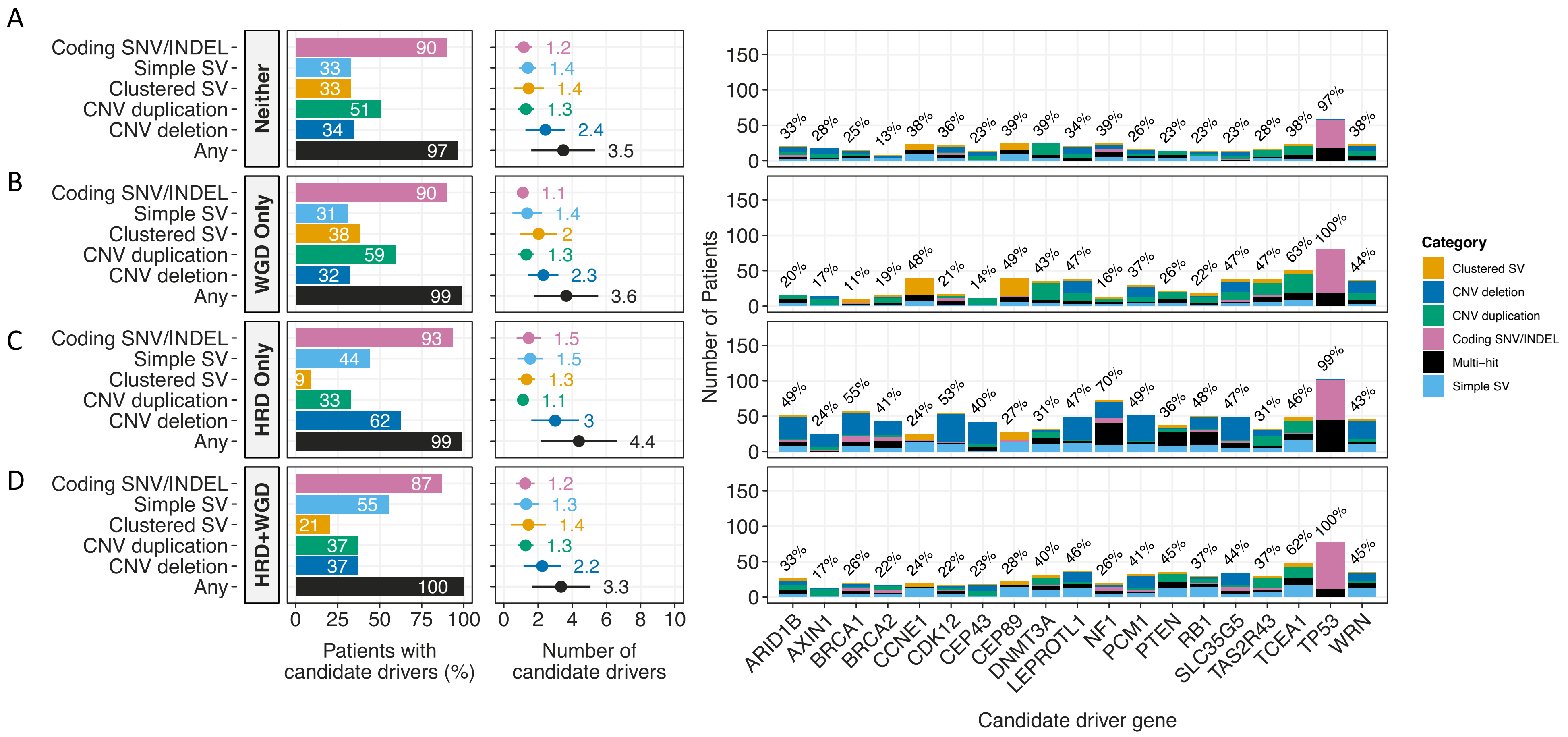


Supp Figure 11: Candidate driver landscape is robust to background genomic states of HRD and WGD. All figures are as described for Figure 3B and C but in a subset of samples. (A) Candidate driver landscape in tumours that are neither HRD nor WGD (N=61). (B) WGD only (N=81). (C) HRD only (N=104) and (D) both HRD and WGD (N=78).

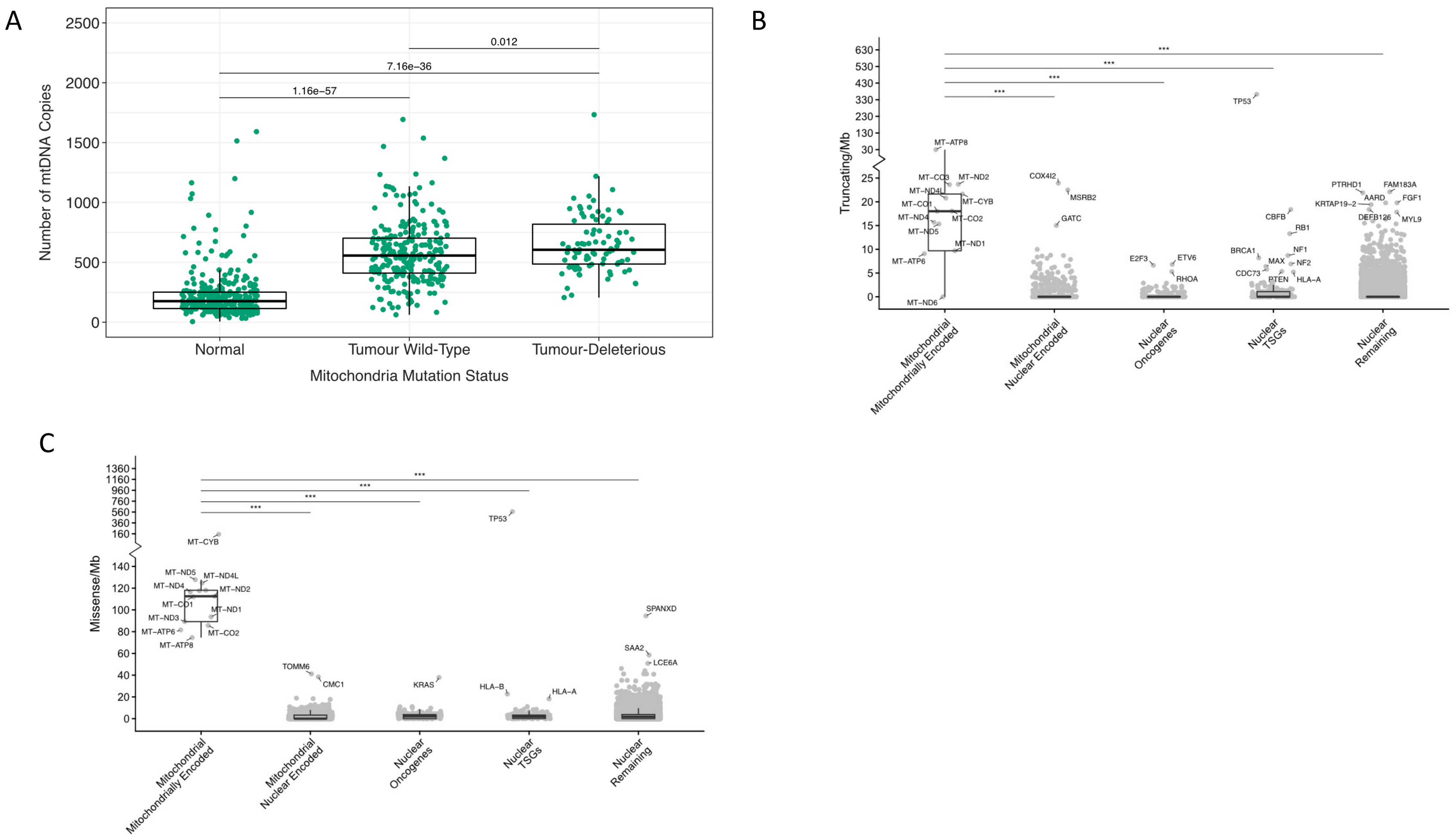


Supp Figure 12: High mitochondrial abundance and somatic mutation loads in HGSOC. (A) Tumour samples have significantly higher mtDNA copy number than normal samples, whether or not tumour samples carry predicted deleterious mtDNA SNVs (deleterious missense variants predicted using SIFT+PolyPhen). Tumour samples show significant ($p<0.001$) excesses of (B) truncating and (C) missense SNVs in mitochondrial genes versus in other gene classes of interest.

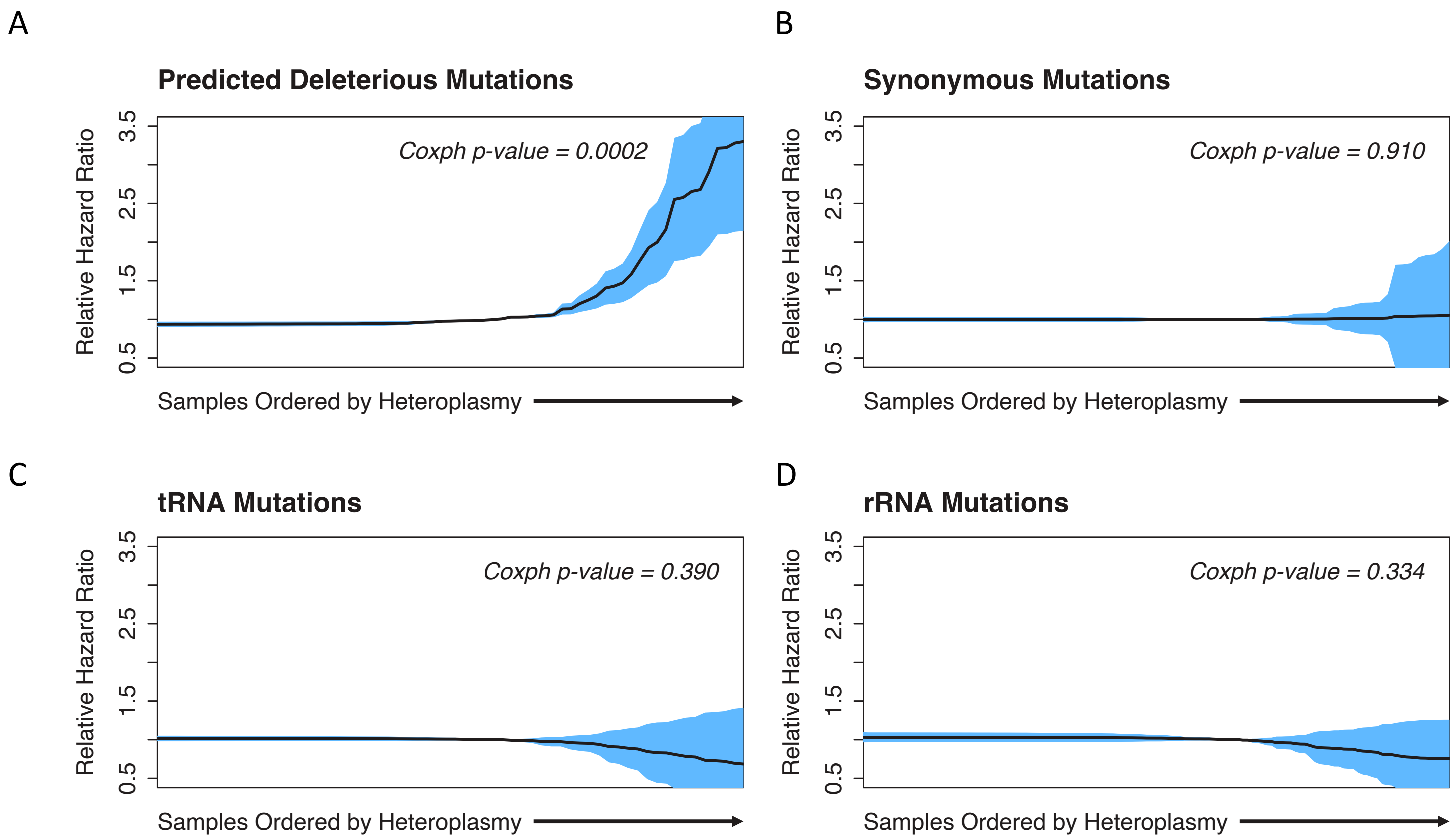


Supp Figure 13: The effects of mtDNA mutation loads on overall survival are mediated by predicted SNV functional impact and heteroplasmy. (A) Cox proportional hazards ratios for predicted deleterious mtDNA SNVs increase for SNVs with higher heteroplasmy. Synonymous SNVs (B), tRNA SNVs (C) and rRNA SNVs (D) are not significantly associated with overall survival at any level of heteroplasmy.

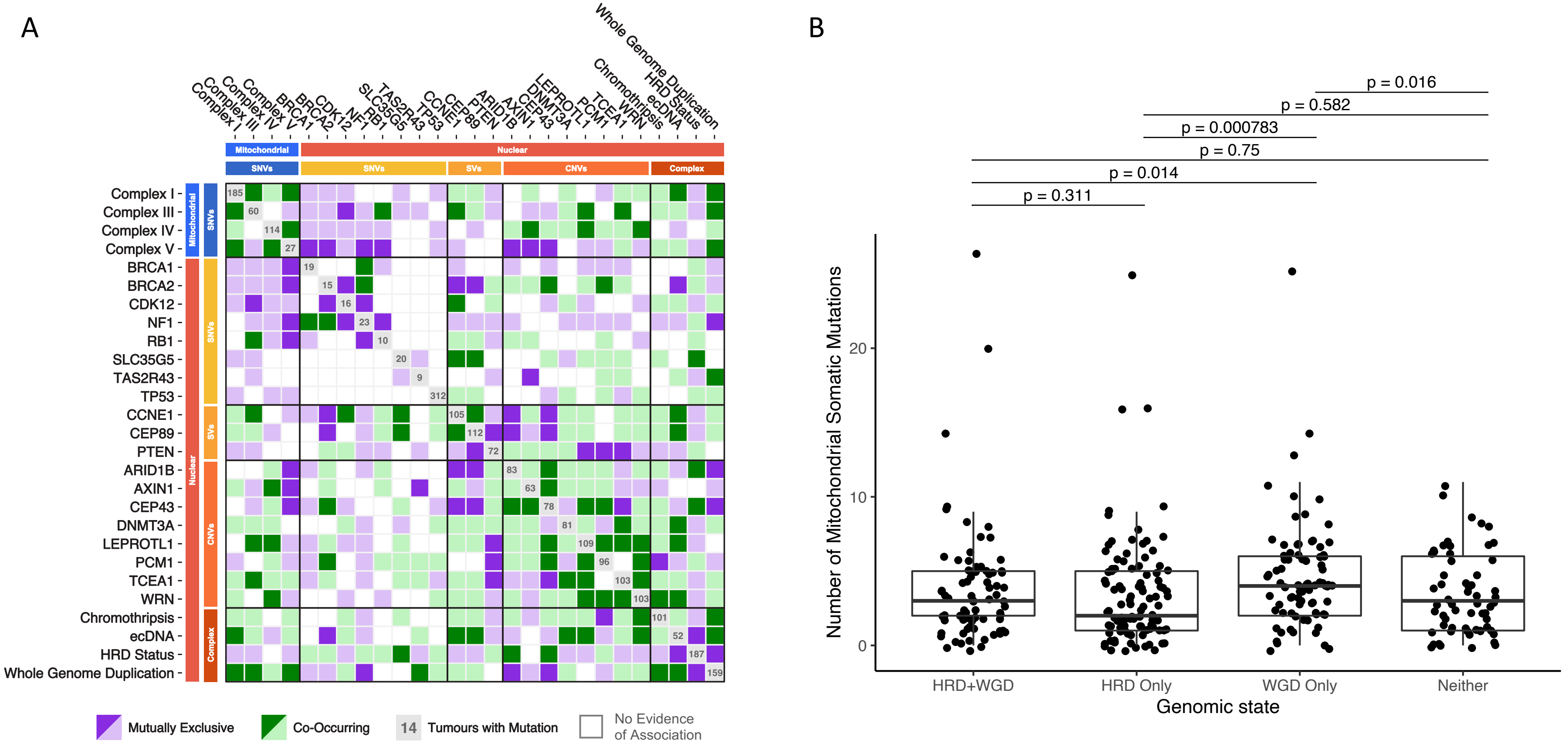


Supp Figure 14: Patterns of co-occurrence and mutual exclusivity between nuclear genome alterations and somatic mtDNA variants. (A) Evolutionary dependencies among nuclear and mtDNA somatic mutations using a Bayesian inference framework (Mina et al, 2020) where significant co-occurrence (green) indicates synergistic interactions and mutually exclusivity (purple) indicates functional redundancies ($p<0.05$). (B) Relative enrichment of mtDNA mutations in samples by background genomic state of HRD and/or WGD.

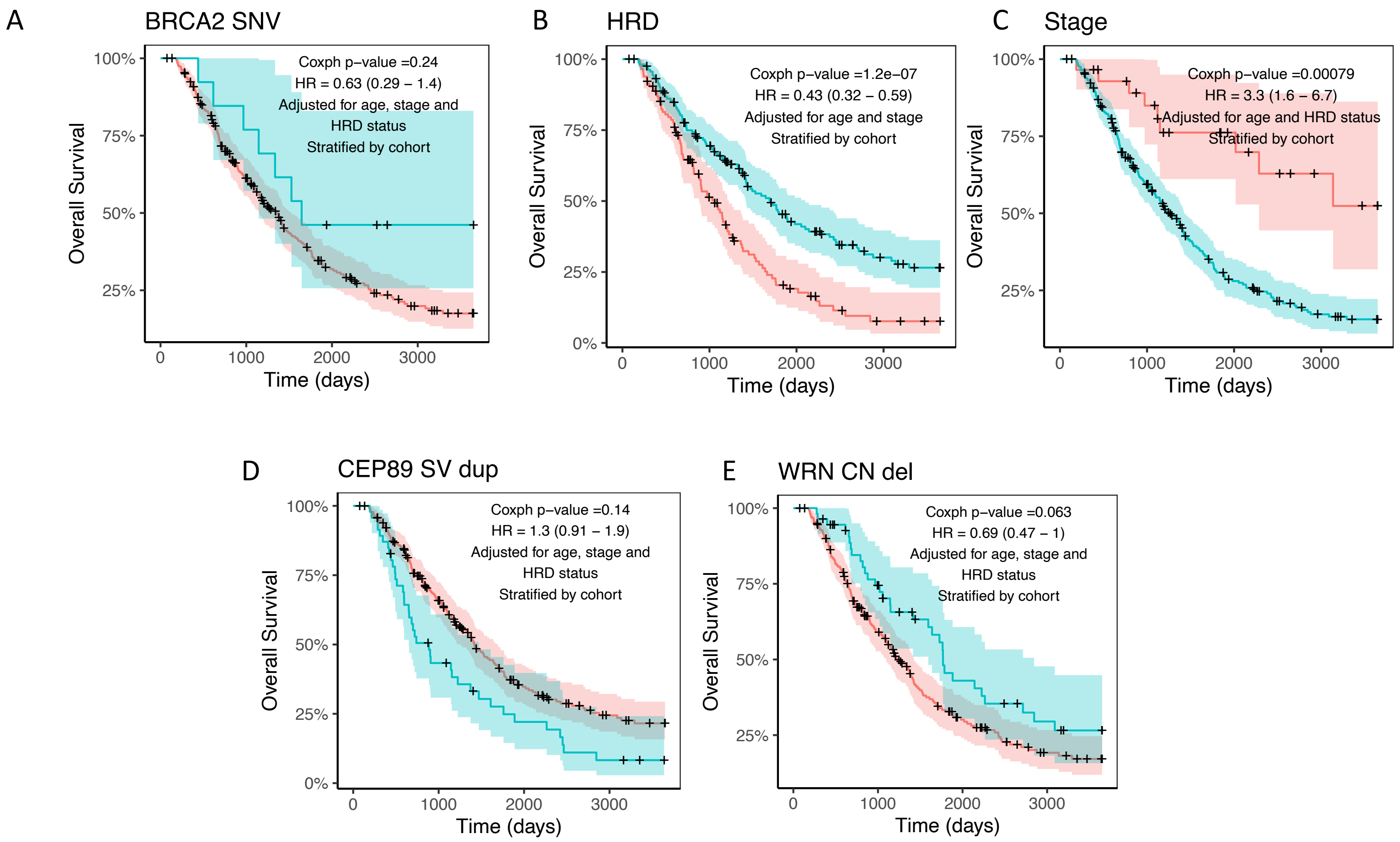


Supp Figure 15: Kaplan-Meier curves of presence (blue curve) and absence (red curve) of selected features (not shown in Figure 6) in the Cox proportional hazards elastic net model of overall survival adjusting for the established effects of age, stage, and HRD status where appropriate and stratifying by cohort. (A) BRCA2 SNV, (B) HRD, (C) stage at diagnosis (D) CEP89 SV duplication and (E) WRN CN deletion. CEP89 SV duplication is highly correlated with CCNE1 SV duplication and likely reflects the same signal as attribution of effects in the presence of high correlation is necessarily arbitrary in the elastic net.
